# Lipoprotein sorting to the cell surface via a crosstalk between the Lpt and Lol pathways during outer membrane biogenesis

**DOI:** 10.1101/2022.12.25.521893

**Authors:** Qingshan Luo, Chengai Wang, Shuai Qiao, Shan Yu, Lianwan Chen, Seonghoon Kim, Kun Wang, Jiangge Zheng, Yong Zhang, Fan Wu, Xiaoguang Lei, Jizhong Lou, Michael Hennig, Wonpil Im, Long Miao, Min Zhou, Yihua Huang

## Abstract

Lipopolysaccharide (LPS) and lipoprotein, two essential components of the outer membrane (OM) in Gram-negative bacteria, play critical roles in bacterial physiology and pathogenicity. LPS translocation to the OM is mediated by LptDE, yet how lipoproteins sort to the cell surface remains elusive. Here we report the identification of an inventory of lipoproteins that are transported to the cell surface via LptDE. Notably, we determined crystal structures of LptDE from *Pseudomonas aeruginosa* and its complex with an endogenous *Escherichia coli* lipoprotein YifL. The *pa*LptDE-YifL structure demonstrates that YifL translocates to the OM via LptDE, in a manner similar to LPS transport. The β-barrel domain serves as a passage for the proteinaceous moiety while its acyl chains are transported outside. Our finding has been corroborated by results from native mass spectrometry, immunofluorescence, and photocrosslinking assays, revealing a unique mechanism through which lipoproteins are translocated across the OM in an ATP- and LPS-dependent manner. Moreover, our study expands the scope of current knowledge of lipoprotein sorting by disclosing a crosstalk between the Lpt and Lol pathways.

## Introduction

Gram-negative bacteria feature a double-membraned cell envelope, of which the outer membrane (OM) serves as a selective permeability barrier. It shields the cells against a wide variety of harmful chemicals while allowing efficient exchange of nutrients and wastes in and out of the cells^1–3^. Structurally, OM is an asymmetrical lipid bilayer with lipopolysaccharides (LPS) enriched in its outer leaflet and phospholipids in its inner leaflet, respectively. In addition, OM harbors two types of proteins, namely integral outer membrane proteins and lipoproteins. The former, also referred to as OMPs are almost exclusively β-barrel structured and effectuate essential functions including OM biogenesis, secretion and efflux pumps^2,4,5^ (Extended Data Fig. 1A). The latter are also called outer membrane lipoproteins that attach to the membrane via its N-terminal lipid tails. While the globular domains of most OM lipoproteins reside within the periplasm, a subset can be detected by probes outside of the cell and accordingly termed as “surface exposed lipoproteins (SLPs)”. Thereby SLPs, together with LPS are the two major lipid-containing structural elements of the OM that are exposed to the extracellular milieu (Extended Data Figs. 1A and 1B), where they can be recognized by Toll-like receptors (TLRs) 4 and 2, respectively, on host cells, eliciting a sequence of events associated with pathogenicity^6–8^.

Synthesized in the cytoplasm and matured in the inner membrane (IM), both LPS and SLPs are amphipathic molecules. In Gram-negative bacteria, trafficking of these molecules across the aqueous periplasm to the OM is carried out by two distinct pathways, namely, the lipopolysaccharide transport (Lpt) and the localization of lipoprotein (Lol) pathways (Extended Data Fig. 1A). Lpt pathway comprises seven essential Lpt proteins (LptA-G)^9,10^. Among them the heteromeric ATP-binding cassette (ABC) transporter LptB_2_FG extracts LPS from the outer leaflet of the IM, and transfers it to the membrane-bound protein LptC. The periplasmic protein LptA connects the LptB_2_FGC complex with the LptDE complex in the OM^9,11–19^. Release of the LPS from LptDE is facilitated via lateral opening at the first and last β strands of the LptD β-barrel^12,20–22^. Efficient transport of LPS to the cell surface thus requires assembly of a transenvelope complex that consists of all seven Lpt proteins and utilizes the energy derived from the ABC transporter LptB_2_FG to drive the unidirectional LPS export^10,23,24^.

In contrast, lipoprotein partition to the OM via the classical Lol pathway does not involve the assembly of a transenvelope complex. Rather, a periplasmic protein LolA shuttles nascent lipoproteins, released from the ABC transporter LolCDE, across the periplasm to the OM-attached lipoprotein LolB. LolB receives the lipoproteins from LolA and inserts them to the inner leaflet of the OM^25–28^ (Extended Data Fig. 1A). Despite the current state of knowledge that lipoproteins retain to the periplasmic sides of either the OM or the IM, a plethora of functionally important lipoproteins has been found to be surface-exposed lipoproteins (SLPs) in Gram-negative bacteria ^26,28–31^. However, how these SLPs reach the cell surface remains enigmatic and posts new questions in understanding lipoprotein transport mechanism.

Here we employed a combination of structural, biochemical and mass spectrometric approaches and established an inventory of lipoproteins that are transported to the cell surface via LptDE. Notably, we determined high-resolution crystal structures of LptDE from *P. aeruginosa* (*pa*LptDE) and its complex with an endogenous *E. coli* lipoprotein YifL. We conclude that LptDE translocates YifL to the OM, in a manner similar to LPS transport. Our findings reveal a unique mechanism in which LptA of the Lpt pathway receives lipoproteins from the LolCDE of the Lol pathway and delivers them to the LptDE en route to cell surface.

## Results

### Identification of a list of lipoproteins that interact with the LptDE complex

During purification of LptDE proteins, we noticed that certain endogenous proteins invariably co-purify with LptDE complexes from both *P. aeruginosa* (*pa*LptDE) and *S. flexneri* (*sf*LptDE) (Figs. 1A and 1B). Proteomic analysis using mass spectrometry identified the co-purified proteins as predominantly YifL and Lpp (the Braun’s lipoprotein), two of most abundant lipoproteins in *E. coli*^29,32^. Further analysis yielded seven additional lipoproteins that co-purify with LptDE. These proteins, present in relatively low abundance in *E. coli* were detected *via* concentration of a low-yield *pa*LptDE sample to *ca*. 20 mg ml^-1^ (Fig. 1B). Therefore, our initial analysis led to a complete inventory of 9 lipoproteins, namely YifL, Lpp,YedD, YbjP, SlyB, putative lipoprotein APQ22800.1, OipY, Slp and Blc, that are enlisted as candidates for interaction partners of LptDE (Extended Data Table 1).

**Fig. 1.**
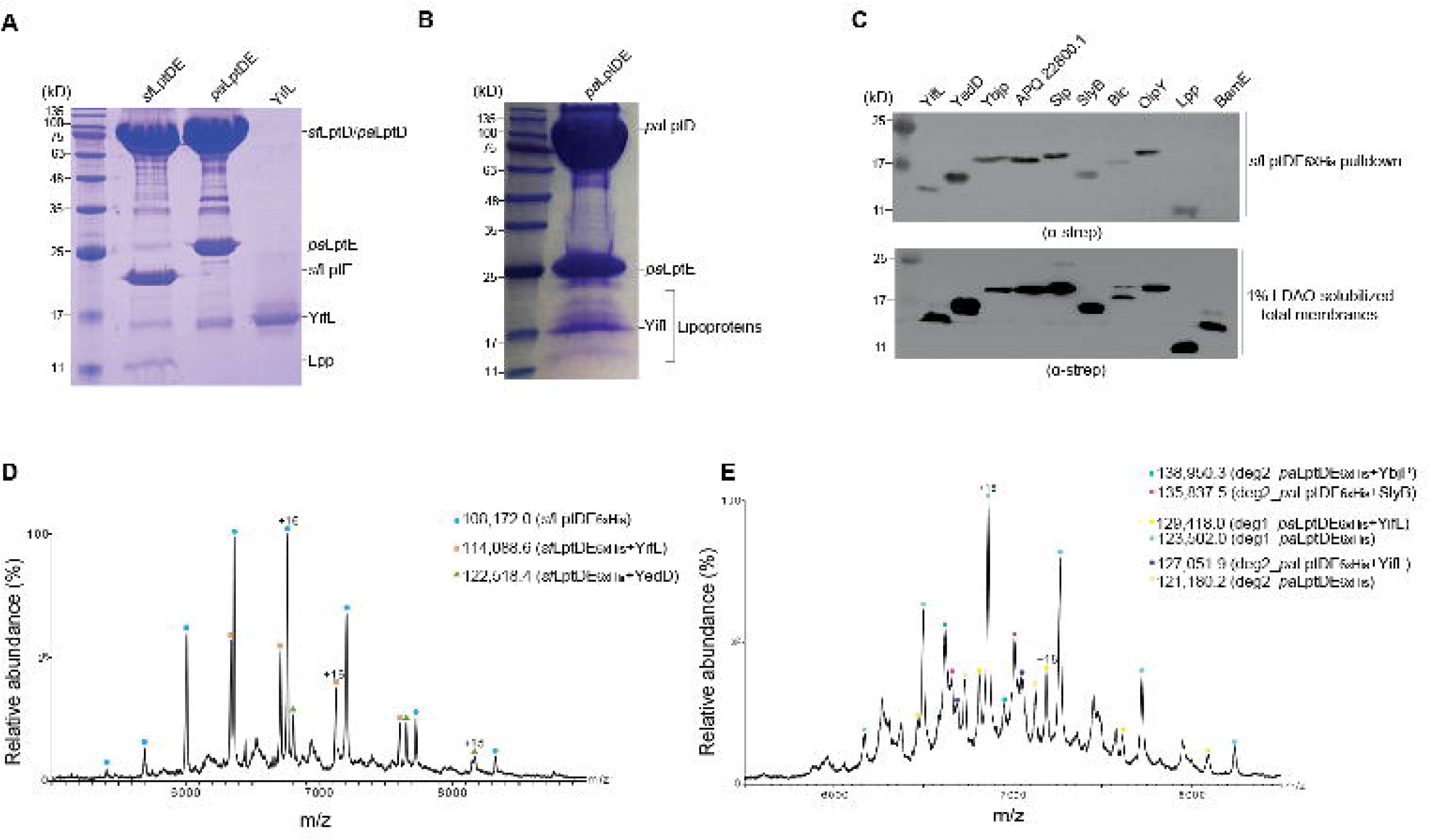
Identification and characterization of an inventory of LptDE-bound lipoproteins. (**A**) SDS-PAGE gel showing that endogenous lipoproteins YifL and/or Lpp were co-purified with affinity-purified LptDE_6×His_ proteins. Endogenous YifL migrated to a similar position to the recombinant YifL_6×his_ on the SDS-PAGE gel (rightmost lane). (**B**) SDS-PAGE analysis of the purified *pa*LptDE_6×His_ protein showing the co-purified endogenous lipoproteins (YifL, Lpp, YedD, YbjP, putative lipoprotein APQ22800.1, SlyB, OipY, Slp and Blc). The lipoproteins were identified by mass spectrometry analysis. The *pa*LptDE_6×His_ sample was concentrated to ~20 mg ml^-1^ and 2 μl sample was loaded. (**C**) Immunoblotting using anti-Strep tag II antibody showing that the affinity-purified *sf*LptDE_6×His_ pulled down all nine lipoproteins identified in (**B**). The bottom panel showing the over-expression of each lipoprotein in the cell by using membrane fractions solubilized with 1% LDAO. Each lipoprotein including BamE with a C-terminal Strep tag II was co-expressed with *sf*LptDE_6×His_ in *E. coli* SF100 strain in the assay. (**D**) Native mass spectrometry analysis of the purified *sf*LptDE_6×His_ protein. The spectrum showing the presence of three protein complexes, *sf*LptDE_6×His_, *sf*LptDE_6×His_-YifL and *sf*LptDE_6×His_-YedD, in the sample. (**E**) Native mass spectrometry analysis of the purified *pa*LptDE_6×His_ protein. The spectrum indicates the presence of six different protein complexes in the sample, *pa*LptD^Δ12^E_6×His_, *pa*LptD^Δ12^E_6×His_-YifL, *pa*LptD^Δ23^E_6×His_, *pa*LptD^Δ23^E_6×His_-YifL, *pa*LptD^Δ23^E_6×His_-YbjP and *pa*LptD^Δ23^E_6×His_-SlyB. *pa*LptD^Δ12^ and *pa*LptD^Δ23^ stand for the N-terminal 12 and 23 residues of the mature *pa*LptD were degraded, respectively. The molecular masses of the mature lipoproteins are listed in Extended Data Table 1, and theoretical and experimentally determined masses of LptDE-lipoprotein complexes are listed in Extended Data Table 2.

We next sought to confirm the interactions between LptDE and the identified lipoproteins. For this, we co-expressed all of the 9 identified lipoproteins together with *sf*LptDE. Pull down assays using six-histamine-tagged *sf*LptDE_6×His_ as a bait protein captured all of the nine lipoproteins that have a Strep-tag II fused to their C-terminus (Fig. 1C, top panel). It is evident from our results that LptDE indeed forms physical interactions with the nine lipoproteins established by proteomics. However, not all lipoproteins are tightly associated with LptDE.

To further confirm the interaction and to investigate the binding specificity and stoichiometry of the identified lipoproteins to LptDE, we subjected the purified *sf*LptDE and *pa*LptDE samples to native MS analysis. Native MS spectra clearly showed that two of the lipoproteins, YifL and YedD formed stable complexes with *sf*LptDE at unit stoichiometry (Fig. 1D), whereas *pa*LptDE exhibited tight association with YifL, SlyB and YbjP, three of the earlier MS-identified lipoproteins (Fig. 1E). These observations reinforced the notion that LptDE selectively binds a subset of lipoproteins with defined stoichiometry.

### Overall structure of the *pa*LptDE-YifL complex

A more convincing evidence of LptDE:lipoprotein interaction comes from crystallographic studies. We therefore sought to obtain an atomic structure of *pa*LptDE-YifL and uncover the interaction details. *Pa*LptD bears a low sequence identity (<25%) with the two LptD homologues with full-length structures to date^21,22,33^, containing an extra 120-residue loop at the N-terminus (Fig. 2A and Extended Data Fig. 2). After extensive crystallization trials, crystals obtained from the purified *pa*LptDE sample diffracted to 3.0-Å resolution. The structure was determined using molecular replacement with the structure of a jellyroll-truncated *pa*LptDE as search model ^22^ and was refined to R_work_/R_free_=0.22/0.26 (Table 1). Throughout model building, we noticed a continuous, strong positive difference Fourier electron density that could not be accounted for by *pa*LptDE itself (Fig. 2B). The ribbon-shaped electron density is clogged at the interface between the β-jellyroll and the β-barrel domains of *pa*LptD: the electron density segment located in the *pa*LptD β-barrel lumen displays characteristic polypeptide features with discernible side chains, whereas the trifurcated branches extending into the membrane boundary are reminiscent of three acyl chains of a lipoprotein (Fig. 2B). We therefore conclude that the extra density corresponds to a molecule of YifL in association with *pa*LptDE. SDS-PAGE analysis of the dissolved crystals confirmed the presence of YifL in the crystals (Extended Data Fig. 3). In the final *pa*LptDE-YifL structure, the N-terminal 81 residues of the mature *pa*LptD and the C-terminal 35 residues of YifL are invisible. We attribute this to intrinsic flexibility as well as partial protein degradation in the aforementioned regions. The latter has been confirmed by Edman sequencing and MS analysis (Fig. 1E and Extended Data Table 2). Overall, our *pa*LptDE-YifL structure resembles a previous β-jellyroll-truncated *pa*LptDE structure with a root-mean-square deviation (RMSD) of 2.0 Å over 740 aligned Cα atoms. Besides, we also observed a long α-helix at the C-terminus of *pa*LptE (Fig. 2B), a feature that is consistent with the NMR structure of the isolated *pa*LptE but missing in the previously-reported truncated *pa*LptDE structure ^34 22^.

**Fig. 2.**
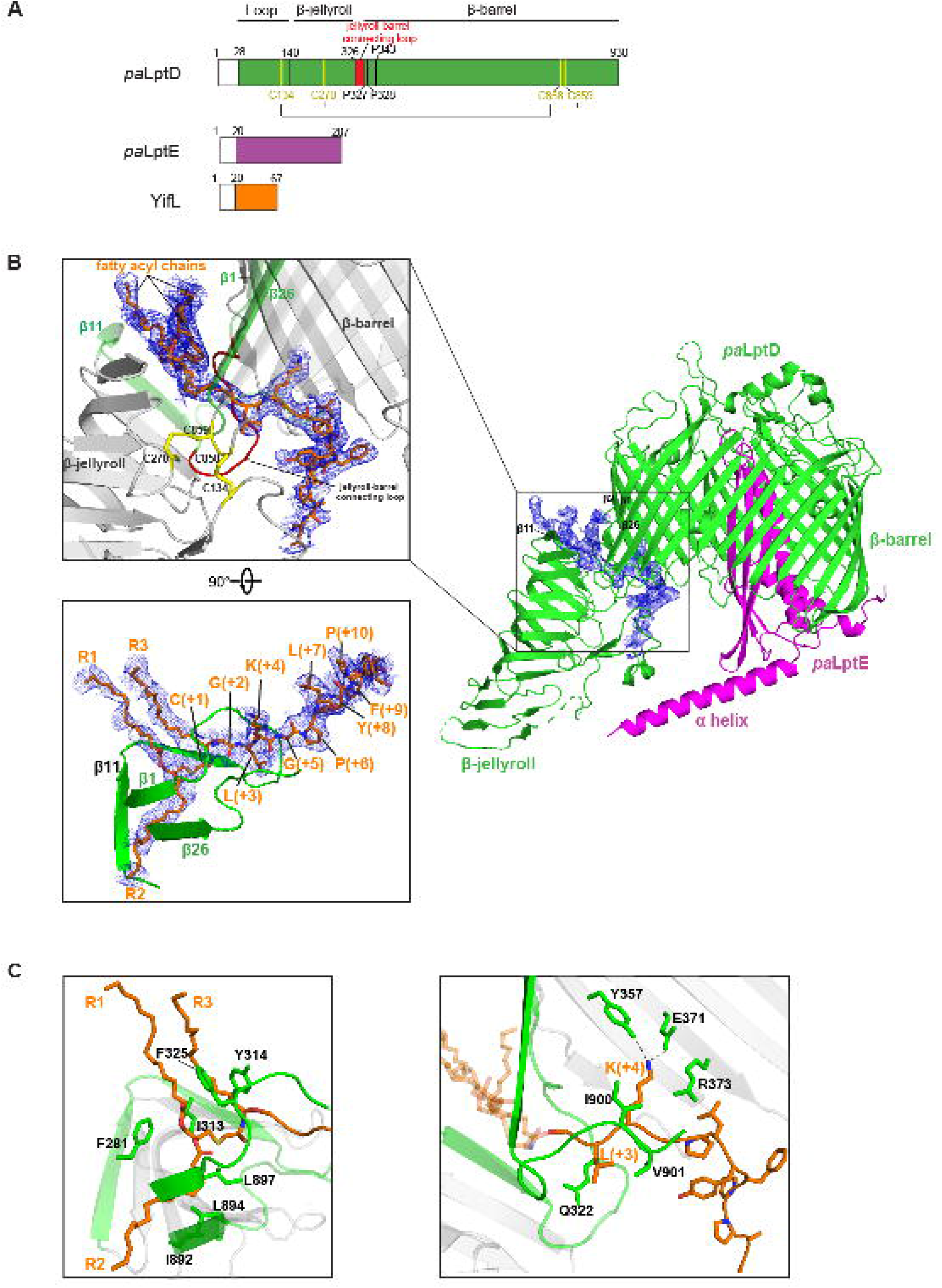
Crystal structure of the *pa*LptDE-YifL complex. (**A**) Schematic structures of *pa*LptD (green), *pa*LptE (magenta) and YifL (orange). The two pairs of conserved inter-domain disulfide bonds (yellow) and the jellyroll-barrel connecting loop (residues P316-P328, red) in *pa*LptD are labeled and highlighted, respectively. The signal peptides of *pa*LptD (residues 1-28), *pa*LptE (residues 1-20) and YifL (residues 1-20) are labeled. (**B**) Unbiased *F_o□_–□F_c_* difference Fourier electron density (blue mesh, contoured at 2.0 σ) calculated before modeling YifL molecule. *Pa*LptD (green) and *pa*LptE (magenta) are shown in cartoon. The YifL (stick mode) was placed in the electron densities, showing that the electron density fragment in the *pa*LptD barrel displays polypeptide features with bulky side chains (left top insert). The jellyroll-barrel connecting loop and the two pairs of disulfide bonds are highlighted in red and magenta, respectively. Numbering of the YifL residues in the structure follows the nomenclature of bacterial lipoproteins (left bottom insert). (**C**) Close-up view of the YifL-*pa*LptDE interactions. Residues of *pa*LptD (F281, I313, Y314, F325, I892, L894 and L897, in green) that interact with the three acyl chains (R1, R2 and R3) are shown in stick mode (left panel); residues of *pa*LptD (Q322, Y357, E371, R373, I900 and V901, in green) that interact with the proteinaceous part of YifL are also shown in stick mode and labeled (right panel). YifL is shown in stick mode (brown). R1, R2 and R3 stand for three acyl chains of YifL.

**Table 1:** Data collection and refinement statistics

|  | *pa*LptDE-YifL | Apo-*pa*LptDE |
| --- | --- | --- |
| **Data collection** |  |  |
| Space group | $P2_1$ | $P2_12_12_1$ |
| Cell dimensions |  |  |
| *a, b, c* (Å) | 93.9, 140.5, 133.2 | 113.0, 160.5, 218.0 |
| $\alpha, \beta, \gamma$ (°) | 90.0, 104.4, 90.0 | 90.0, 90.0, 90.0 |
| Wavelength (Å) | 0.9792 | 0.9792 |
| Resolution (Å) | 50-2.98<br>(3.09-2.98) | 50-3.30<br>(3.44-3.30) |
| $R_{\text{sym}}$ | 0.12(1.0) | 0.16(1.0) |
| $I/\sigma$ | 12.7(1.7) | 10.8(1.7) |
| CC<sub>1/2</sub> (%) | 97.0(57.3) | 96.0(36.9) |
| Redundancy | 5.0 (5.1) | 4.0(3.7) |
| Completeness (%) | 97.2(77.6) | 96.4(94.0) |
| **Refinement** |  |  |
| Resolution (Å) | 2.98 | 3.30 |
| No. reflections | 66447 | 59014 |
| $R_{\text{work}}/R_{\text{free}}$ | 23.4/26.4 | 22.5/26.3 |
| No. atoms | 15664 | 15855 |
| B-factors | 80.91 | 80.81 |
| R.m.s. deviations |  |  |
| Bond lengths (Å) | 0.010 | 0.010 |
| Bond angle (°) | 1.27 | 1.09 |
| Ramachandran plot |  |  |
| Favoured (%) | 93.6 | 95.0 |
| Allowed (%) | 5.8 | 4.6 |
| Outliers (%) | 0.6 | 0.4 |

\*The numbers in parentheses are for the highest resolution shell

Importantly, our structure shows that three hydrophobic acyl chains of YifL are positioned in a hydrophobic environment surrounded by the last V-shaped β strand of the β-jellyroll and the exterior surface of the *pa*LptD β-barrel. This suggests that the hydrophobic acyl chains of YifL directly entered the OM upon leaving the *pa*LptD β-jellyroll (Fig. 2B). In contrast, the proteinaceous moiety of the mature YifL (residues Gly2-Tyr8) interacts with residues lining the hydrophilic lumen of the LptD β-barrel (Fig. 2C, right panel). This combined with the periplasm-resided C-terminus, indicates that the proteinaceous moiety of YifL will eventually tuck into the hydrophilic lumen of the LptD β-barrel. Remarkably, YifL is clogged in a region that is covalently sealed by the jellyroll-barrel connecting loop at one side and two conserved inter-domain disulfide bonds at the opposite side (Fig. 2B, left top panel). Hence, release of YifL from *pa*LptD β-barrel to the OM can only proceed through unzipping the non-covalent interface between β1 and β26 of the *pa*LptD β barrel ^21^. Taken together, the *pa*LptDE-YifL structure points to a unique mechanism through which amphipathic lipoproteins are transported to the cell surface by partitioning their hydrophobic acyl chains and hydrophilic moiety into the OM space and hydrophilic lumen of *pa*LptD, respectively, in a manner similar to LPS translocation. The structure also highlights a critical role of the two conserved bridging disulfide bonds in preventing lipoproteins from mistakenly entering the inner leaflet of the OM during the export process^35^.

### Conformational changes of the β-jellyroll domain of *pa*LptD upon YifL binding

To explore conformational changes of *pa*LptDE upon YifL binding, we sought to obtain the apo-*pa*LptDE structure and purified the recombinant *pa*LptDE protein from the *yifL*-depleted *E coli* SF100 strain. The structure was determined to a resolution of 3.3 Å (Table 1). The crystals belong to space group P2_1_2_1_2_1_ with two apo-*pa*LptDE complexes (*pa*LptDE_A and *pa*LptDE_B) in one asymmetric unit. We note that there is only one pair of disulfide bond (Cys134-Cys858) formed in each of the apo-*pa*LptDE complex (Fig. 3A). Intriguingly, for the two *pa*LptDE complexes within one asymmetric unit the individual domain structures (β-jellyroll, β-barrel and LptE) are almost identical. The two β-jellyroll domains however, show a 6° rotation along the axis perpendicular to the membrane plane (Fig. 3A). This suggests flexibility of the β-jellyroll domain with only one disulfide link, implying that both pairs of conserved inter-domain disulfide bonds within *pa*LptD are required to maintain the orientation of the β-jellyroll ^35^.

**Fig. 3.**
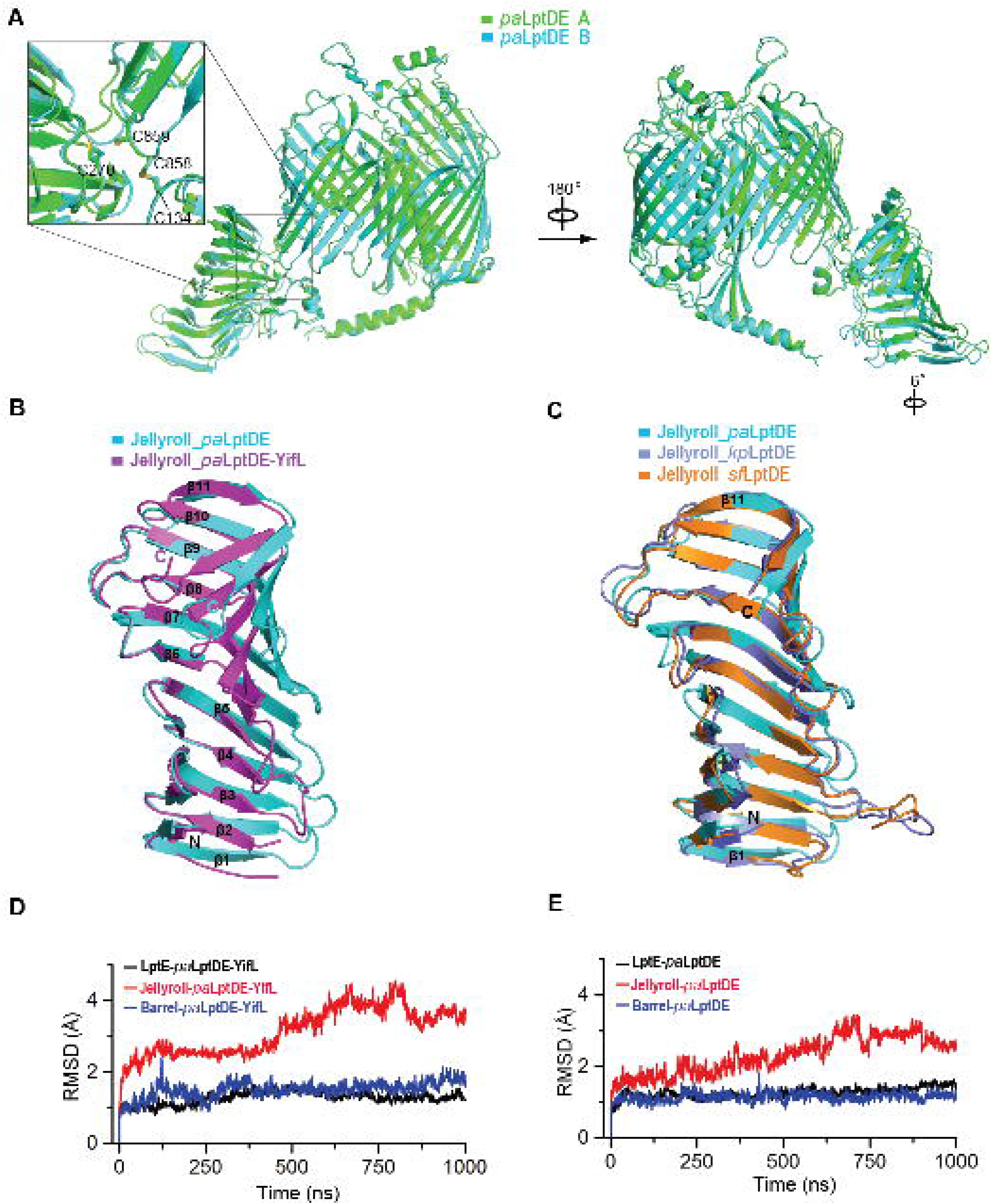
Structural comparison and analysis of the β-jellyroll domain in apo-*pa*LptD and in *pa*LptD-YifL. (**A**) Structural overlay of the two apo-*pa*LptDE complexes (*pa*LptDE_A and *pa*LptDE_B) in the asymmetric unit showing that there is only one pair of disulfide bond (C134-C858) formed in each complex (left) and a 6° rotation of the β-jellyroll domain (right). (**B**) Superimposition of the β-jellyroll domains of *pa*LptDE-YifL (violet) with that of apo-*pa*LptDE (cyan). The β-jellyroll domain consists of 11 β strands that are numbered from the N-terminus of the domain. The β-jellyroll domain in *pa*LptDE-YifL adopts a closed conformation as compared to that in apo-*pa*LptDE with a shortened distance of the two ends for each of the 11 β strands, ranging from 0.5 Å to 4.3 Å (Extended Data Table 3). (**C**) Structural overlay of three β-jellyroll domains of apo-LptDE homologue structures showing the β-jellyroll domains all adopt an open conformation: *pa*LptDE (cyan), *kp*LptDE (slate; PDB code: 5IV9) and *sf*LptDE (brown; PDB code:4Q35). *kp*LptDE stands for the LptDE homolog from *klebsiella pneumoniae.* (**D-E**) Average backbone RMSDs of the simulated structures (*pa*LptE, β-jellyroll and β-barrel) in *pa*LptDE-YifL (**D**) and apo-*pa*LptDE (**E**) structures showing the dynamic nature of the β-jellyroll domain in both structures.

Structural overlay of apo-*pa*LptDE with *pa*LptDE-YifL reveals that YifL binding not only caused a slightly enlarged β-barrel lumen on the periplasmic side and a remarkable rearrangement of the jellyroll-barrel connecting loop (Extended Data Figs. 4A and 4B), but also induced the β-jellyroll to adopt an overall closed conformation (Fig. 3B). Specifically, in comparison to apo-*pa*LptDE, the β-jellyroll domain of *pa*LptDE-YifL has reduced distances between the ends of each V-shaped β strands. The reductions are in the range from 0.5 Å to 4.3 Å (Extended Data Table 3). Overlay of the β-jellyroll domain of apo-*pa*LptDE with the two existing structures of LptDE homologue further supports this observation ^21,22^ (Fig. 3C). Considering that YifL only makes marginal hydrophobic contacts with the residues from the last V-shaped β strand of the β-jellyroll (Fig. 2C, right panel), the substantial overall conformational change of the jellyroll is most likely attributable to its dynamic nature. In agreement with this, the average B factors of Cα atoms in the β-jellyrolls in both *pa*LptDE and *pa*LptDE-YifL structures are higher than their respective β-barrel domain and LptE (Extended Data Figs. 5A and 5B). To corroborate this, we carried out molecular dynamics (MD) simulations in an *E. coli* OM environment and the results showed that the RMSDs of the β-jellyroll domains in both apo- *pa*LptDE and *pa*LptDE-YifL are higher than those of the β-barrel domain and *pa*LptE (Fig. 3D and 3E). Collectively, our studies suggest that the β-jellyroll domain of *pa*LptD is dynamic and its conformation may fluctuate in the process of lipoprotein or LPS export. We speculate that the dynamic nature and the concerted conformational changes may be ubiquitous in all the jellyroll-containing Lpt proteins with which the whole β-jellyroll bridge couples the chemical energy derived from cytoplasmic ATP hydrolysis with mechanical force to drive the export of lipoproteins or LPS molecules across the periplasm.

### LptDE-bound lipoproteins are surface-exposed

The *pa*LptDE-YifL structure implies that lipoprotein YifL, like LPS, will be eventually transported to the cell surface through LptDE (Fig. 2B). To test whether YifL and the other LptDE-bound lipoproteins are indeed surface-exposed, we carried out immunofluorescence assays. For this, each of the identified lipoproteins with a C-terminal His tag was cloned into pET22b vector which was transformed into *E. coli* SF100 cells. We then incubated the resultant *E. coli* cells with primary anti-His antibody, followed by Alexa Fluor 546-conjugated (red-fluorescing) secondary antibody for fluorescence detection. As shown in Fig. 4A, *E. coli* cells that harbored each of the lipoprotein-expressing plasmids displayed red fluorescence circles, suggesting that the C-termini of these lipoproteins are surface-exposed. By contrast, *E. coli* cells transformed with a plasmid that expressed BamB_6×His_, a lipoprotein of the β-barrel-assembly machinery (BAM) complex that attaches to the inner leaflet of the OM ^36,37^, displayed red fluorescence only when the cell membranes were permeabilized (Fig. 4A). Consistent with immunofluorescence assay results, dot blot assays also showed that all the LptDE-bound lipoproteins but not BamB_6×His_ were detected on the intact cells (Figs. 4B). However, it showed that not 100% of the YifL molecules were transported to the cell surface (Figs. 4B), a scenario that echoes a previous study of Lpp^29^. Taken together, our results confirmed that a certain percentage of each of the identified LptDE-binding lipoproteins can be transported to the cell surface.

**Fig. 4.**
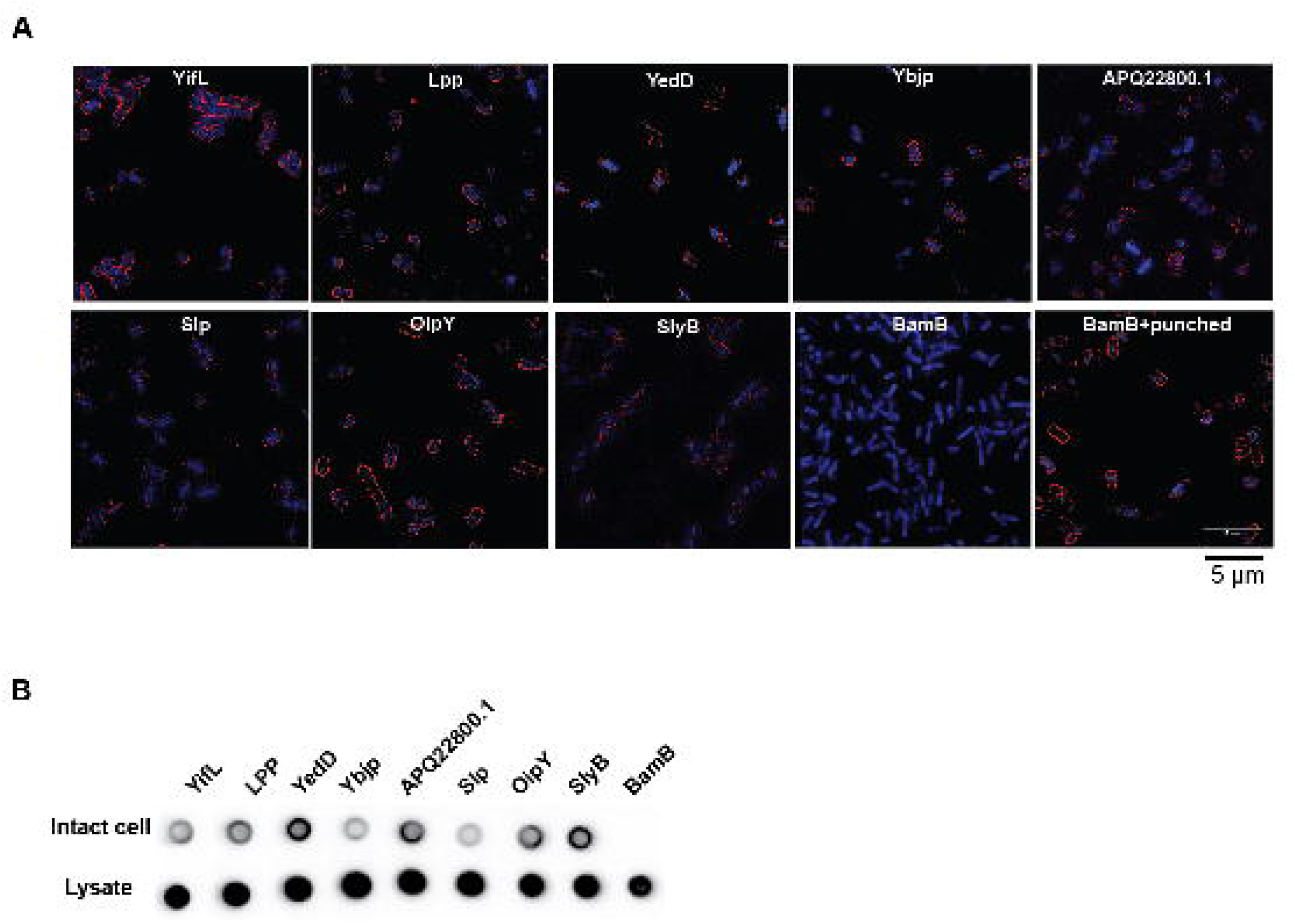
The identified LptDE-bound lipoproteins are exposed on the surface of *E. coli.* (**A**) Immunofluorescence assays showing that YifL, Lpp, YedD, YbjP, APQ22800.1, Slp, OipY and SlyB are surface-exposed. DAPI and Alexa Fluor 546-conjugated secondary antibody are blue- and red-fluorescencing, respectively. BamB was selected as negative control. Membrane permeabilization was carried out by treatment with 0.5% Triton X-100 and 5 mM EDTA. (**B**) Dot blot assays (both using intact cells and cell lysates) showing that the identified LptDE-bound lipoproteins are surface-exposed at least for certain percentage of the expressed lipoproteins in the cells. BamB was selected as negative control. All the lipoproteins contained a C-terminal 6×His tag for visualization or detection.

### YifL is extracted from the IM by LolCDE rather LptB_2_FG

Next, we ask how lipoprotein YifL is transferred to LptDE. As both matured lipoproteins and LPS anchor to the outer leaflet of the IM via their acyl chains prior to trafficking, two potential routes are possible for lipoproteins sorting to the cell surface. First, YifL might be promiscuously extracted by LptB_2_FG from the IM and are transported to the cell surface via the Lpt pathway, given that both lipoproteins and LPS share hydrophobic acyl chains and hydrophilic components (Extended Data Fig. 1B). The second possibility is that YifL, like conventional lipoproteins, is extracted by LolCDE in the Lol pathway but is delivered to LptA, the periplasmic component that bridges the periplasmic gap between LptB_2_FGC and LptDE in the Lpt pathway. To clarify the YifL sorting route, we first probed whether YifL passes through the β-jellyroll domain of LptD. To this end, a photocrosslinkable unnatural amino acid, *p*Bpa, was introduced at either I230 or Y112, two LPS crosslinkable sites in LptD^38,39^. Photocrosslinking were performed when either LptD^I230*p*Bpa^E or LptD^Y112*p*Bpa^E was co-expressed with YifL^23^. We observed YifL cross-linked to LptD at each site (Fig. 5A), suggesting that YifL was delivered to the jellyroll-barrel interface as what we observed in the structure through the β-jellyroll domain of LptD.

**Fig. 5.**
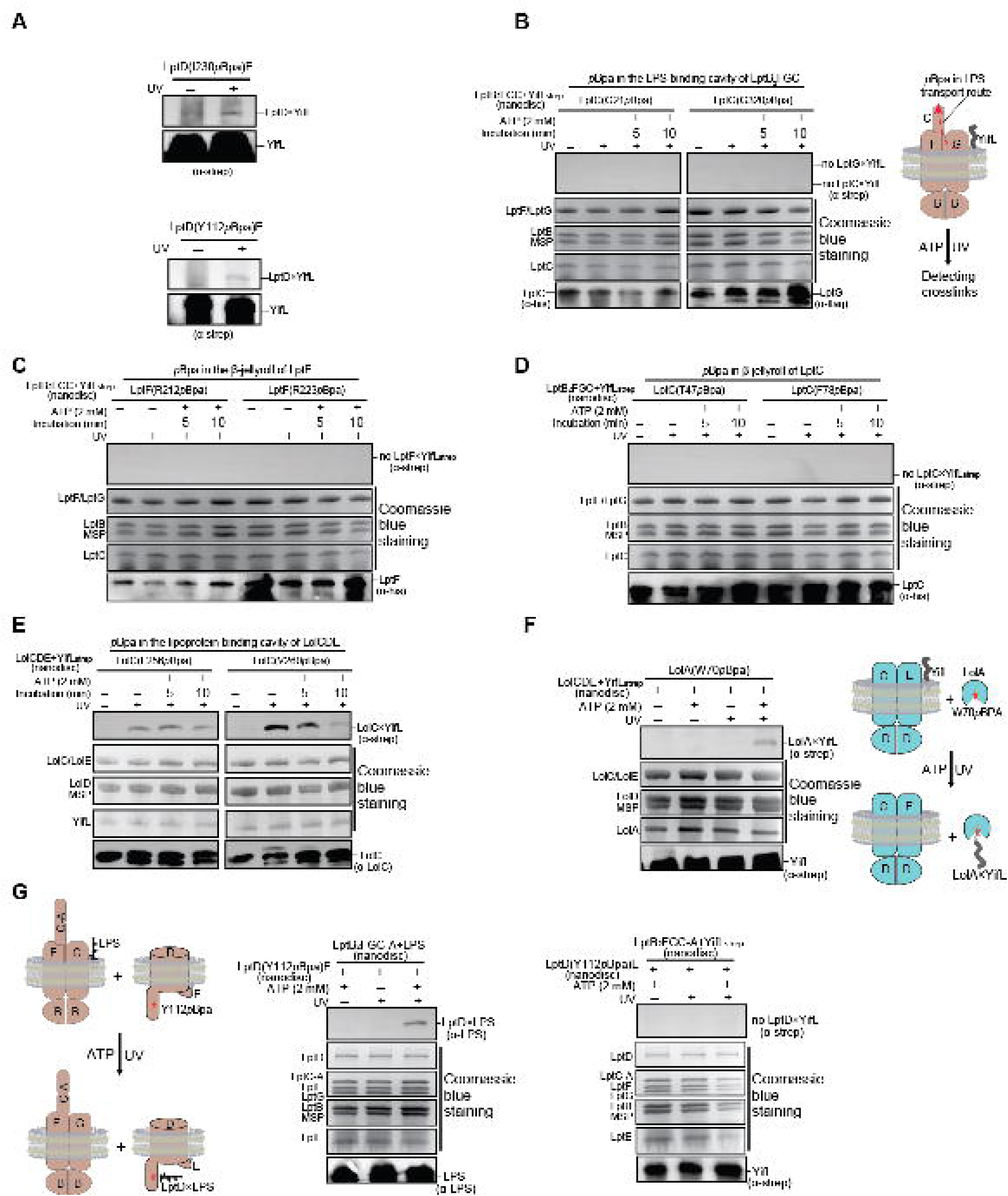
YifL is extracted from the IM by LolCDE instead of LptB_2_FG. (**A**) *In vivo* photocrosslinking showing that YifL cross-linked to *p*Bpa-containing LptD (I230*p*Bpa or Y112*p*Bpa) when YifL and *sf*LptDE were co-expressed in *E. coli.* YifL contained a C-terminal Strep tag II for immunoblotting detection. (**B-D**) Reconstitution of YifL membrane-to-membrane transport. Residues G21 of LptC and G320 of LptG are located in the LPS-binding cavity of LptB_2_FGC, but neither LptC (G21*p*Bpa) nor LptG (G320*p*Bpa) cross-linked to YifL (**B**) Residues R212 and R223 of LptF are located in the β-jellyroll of LptF, but neither LptF (R212*p*Bpa) nor LptF (R223*p*Bpa) cross-linked to YifL (**C**). Residues T47 and F78 of LptC are located in the β-jellyroll of LptC. Neither LptC (T47*p*Bpa) nor LptC (F78*p*Bpa) cross-linked to YifL (**D**). Cartoons show experimental designs of the reconstituted system. YifL can be inserted into nanodisc in either orientation, but only the productive orientation is shown for simplicity. The red arrow denotes LPS transport direction in LptB_2_FGC. None of these selected sites cross-linked to YifL, with or without ATP, but all of these selected sites in LptB_2_FGC cross-linked to LPS upon being substituted with *p*Bpa (Extended Data Figs. 6A-C). (**E**) YifL cross-linked to LolC in LolCDE upon UV radiation. Residues L256 and V269 of LolC are located in the lipoprotein-binding cavity of LolCDE. The YifL×LolC crosslinks decreased with the increase of incubation time. (**F**) LolA cross-linked YifL upon being released from LolCDE in an ATP-dependent manner. Residue W70 of LolA was substituted with *p*Bpa in the assay. Cartoons show experimental designs of the reconstituted system. The red star denotes the position of *p*Bpa in LolA. (**G**) *In vitro* LPS or YifL membrane-to-membrane transport assay showing that LPS was efficiently transported from LptB_2_FGC-A to LptD^Y112*p*Bpa^E in an ATP-dependent manner (middle), but YifL was not transported from LptB_2_FGC-A to LptD^Y112*p*Bpa^E (right). Cartoons show experimental designs of the reconstituted system.

To investigate whether YifL is extracted by LptB_2_FGC from the IM, we sought to reconstitute YifL transport using purified protein components. First, to monitor whether YifL enters into the LPS-binding cavity of LptB_2_FGC, two LPS-photocrosslinkable sites (LptC^G21^ and LptG^G320^) in the LPS-binding cavity of LptB_2_FGC were substituted with *p*Bpa. The purified LptB_2_FGC^G21*p*Bpa^ or LptB_2_FG^G320*p*Bpa^C complexes were reconstituted in nanodiscs together with either LPS or YifL. We observed that LPS cross-linked to LptC and LptG (Extended Data Fig. 6A). However, YifL did not cross-link to LptC or LptG under the same conditions (Fig. 5B). Furthermore, LPS×LptC (or LPS×LptF) crosslinks (Extended Data Fig. 6B and 6C) were detected at each site in the β-jellyroll bridge of LptB_2_FGC (LptF^R212^, LptF^R223^, LptC^T47^ and LptC^F78^) that was substituted with *p*Bpa in an ATP-dependent manner, but none of these sites cross-linked to YifL (Figs. 5C and 5D). These observations are consistent with previous reports that LptB_2_FGC is responsible for extracting and transporting LPS, thus arguing against YifL entering the LPS-binding pocket of LptB_2_FGC and passing through β-jellyroll domains of LptF and LptC.

We next explored whether YifL is extracted by the ABC transporter LolCDE in the Lol pathway. To test this, we also carried out photocrosslinking assays using purified protein components. Based on the LolCDE structures ^40,41^, we purified LolCDE with *p*Bpa substituted at either LolC^L256^ or LolC^V260^, two sites in the lipoprotein-binding cavity of LolCDE. We detected YifL×LolC^L256pBpa^ and YifL×LolC^V260pBpa^ crosslinks upon UV radiation in the absence of ATP (Fig. 5E), suggesting that YifL entered the lipoprotein-binding pocket of LolCDE. To test whether LolCDE delivers YifL to LolA, we purified LolA that contains *p*Bpa at LolA^W70^ and incubated LolA^W70^ with nanodisc-embedded LolCDE and YifL. We observed YifL×LolA crosslinks only in the presence of ATP (Fig. 5F). Combined our results suggest that YifL, like other lipoproteins, is extracted from the IM by LolCDE, followed by delivery to LolA once released from LolCDE. Our observations are also consistent with the results of our earlier dot blot assays in which we observed that a significant amount of YifL molecules attach to the inner leaflet of the OM.

To further confirm that YifL is not transferred to LptDE from LptB_2_FGC, we performed membrane-to-membrane lipoprotein transfer assay using purified protein components. For this, the nanodisc-embedded LptD^I230*p*Bpa^E (a site in LptD that cross-links to both LPS and YifL) was incubated with either nanodisc-embedded LptB_2_FGC-A+YifL or LptB_2_FGC-A+LPS. LPS strongly cross-linked to LptD in the presence of ATP, but we did not observe any crosslinks between YifL and LptD (Fig. 5G). We therefore concluded that it is not likely that LptB_2_FGC extracts YifL from the IM. Rather a crosstalk between the Lpt and the Lol pathways for YifL surface localization must exist, which possibly involves LolCDE.

### LptA crosstalks to the Lol pathway in YifL surface localization

To rule out the possibility that LolA participates in the delivery of YifL to LptDE, we co-expressed *sf*LptDE and YifL in the *lolA*-depleted MG3324 (NR754 *Δlpp ΔrcsB zii::*Tn10 *cpxA24 ΔlolA) E. coli* strain ^42^ and tested whether YifL is transferred to *sf*LptDE *in vivo.* SDS-PAGE analysis of the affinity-purified sample showed that YifL formed a stable complex with *sf*LptDE (Fig. 6A). The result suggests that LolA is not required for YifL transfer to LptDE. Considering the challenge of the amphipathic lipoprotein traversing the periplasm space, there must exist another chaperone that shuttles YifL to LptDE.

**Fig. 6.**
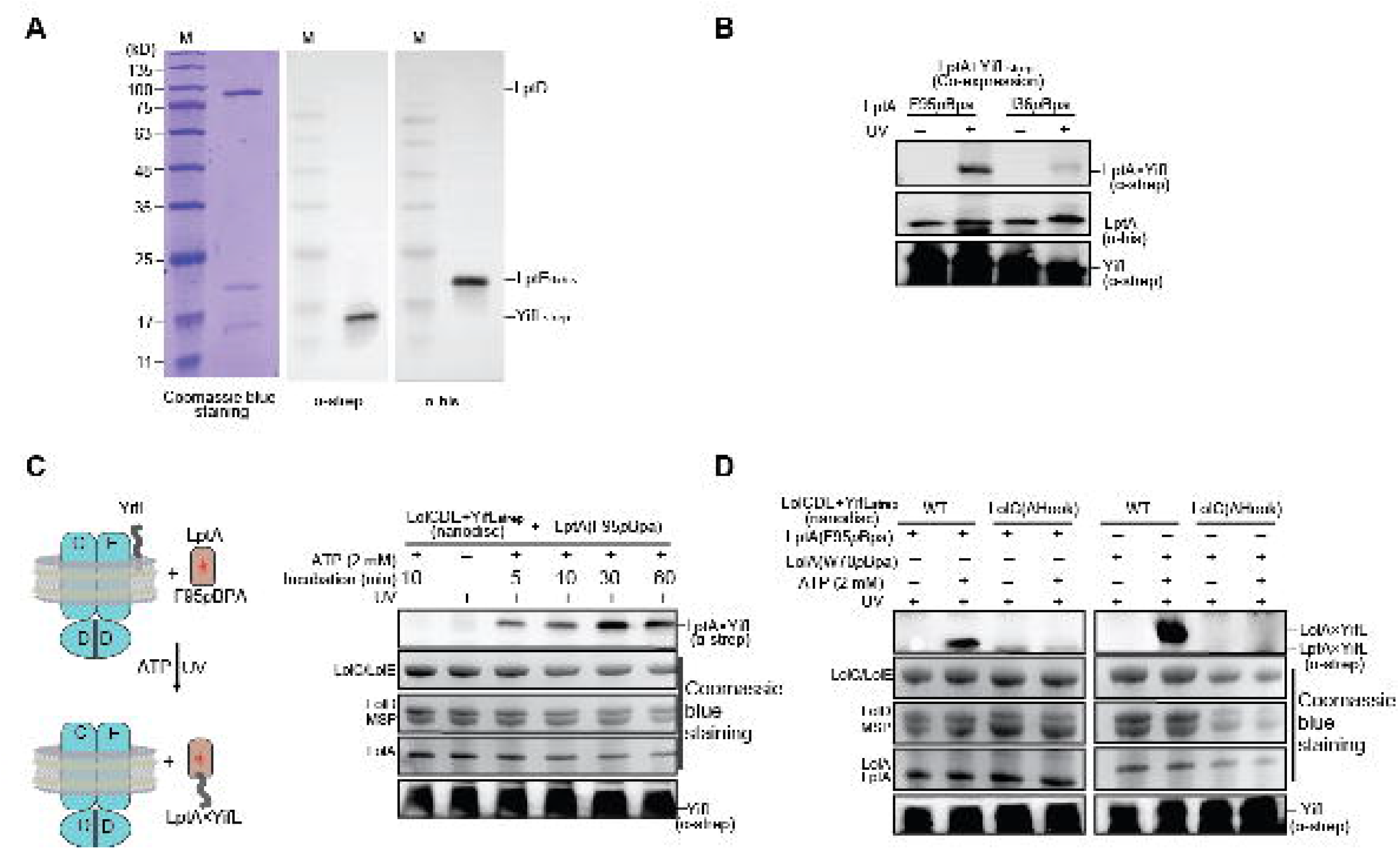
LptA crosstalks to the Lol pathways during YifL surface localization. (**A**) SDS-PAGE (left) and Western blotting (middle and right) analysis of the affinity-purified LptDE_6×His_ when both LptDE_6×His_ and YifL_Strep_ were co-expressed in the *lolA*-depleted MG3324 *E. coli* strain. YifL was copurified with LptDE_6×His_ in the absence of LolA, indicating that LolA is not required for YifL transport to LptDE. (**B**) *In vivo* photocrosslinking assay showing that YifL cross-linked to *p*BpA-containing LptA (F95*p*BpA or I36*p*BpA) when YifL and LptA were co-expressed in *E. coli.* (**C**) *In vitro* YifL transport assay showing that *p*Bpa-containing LptA (F95*p*Bpa) cross-linked to YifL that was reconstituted into nanodiscs together with LolCDE in a time- and ATP-dependent manner. The red star denotes the location of F95*p*Bpa in the LptA β-jellyroll. Cartoons show experimental designs of the lipoprotein transfer assay. (**D**) The Hook motif of LolC in LolCDE is required for YifL transfer from LolCDE to both LolA and LptA.

LptA is the only periplasmic protein that directly interacts with both LptDE and LptB_2_FGC in the Lpt pathway, we examined whether LptA receives YifL upon its release from LolCDE. For this we first co-expressed YifL with LptA that contains *p*Bpa at LptA^F95^ or LptA^I36^ in *E. coli.* We observed that YifL cross-linked to LptA upon UV radiation (Fig. 6B). By contrast, LolB and NlpA, two lipoproteins that respectively anchor to the inner leaflet of the OM and outer leaflet of the IM did not cross-link to LptA in the same scenario (Extended Data Fig. 6D). We further performed *in vitro* lipoprotein transfer assay using purified protein components. We incubated YifL with purified LptA^F95*p*Bpa^ in the presence of nanodisc-embedded LolCDE for photocrosslinking assay. We observed that LptA cross-linked to YifL in an ATP-dependent manner (Fig. 6C). However, when LolCDE was replaced by a variant which lacks the Hook motif (residues P167-P179) in LolC, LptA^F95*p*Bpa^ did not cross-link YifL (Fig. 6D) ^43^. As the Hook motif is critical for lipoprotein transfer to LolA, it is likely that LptA and LolA interact with LolCDE in a similar manner. These findings, together with our earlier lipoprotein transfer assay results, consistently show that LptA crosstalks to the Lol pathway by shuttling YifL across the periplasm and presenting it to LptDE.

### YifL translocates across the OM in an ATP- and LPS-dependent manner

Our earlier studies establish that the LptB_2_FGC complex is not responsible for shuttling YifL over the periplasm. We inquire whether LptB_2_FGC takes part in YifL transport to the cell surface. First, we reconstituted membrane-to-membrane delivery of LPS by incorporating purified inner and outer membrane transport complexes into separate nanodiscs. We found that the purified LptA protein is prone to form oligomers, hampering its binding with LptB_2_FGC or LptDE with high affinity. In order to increase LPS transport efficiency *in vitro,* we constructed an LptC-LptA fusion protein that formed a stable complex with LptB_2_FG. After mixing the nanodisc-embedded LptD^Y112*p*Bpa^E protein complexes with nanodisc-embedded LptB_2_FGC-A+LPS, we observed that LPS cross-linked to LptD in an ATP-dependent manner (Extended Data Fig.7). By contrast, in the absence of LptA, we did not observe any LPS×LptD crosslinks (Extended Data Fig. 7.). The results demonstrate the *in vitro* reconstitution of Lpt pathway in which all seven Lpt proteins and ATP are required for LPS transport. Next, we investigate whether reconstituted Lpt pathway drives the export of LptDE retained YifL across the OM. We then prepared three different types of nanodisc-embedded protein complex samples (LptB_2_FGC-A, LptB_2_FGC-A+YifL and LptB_2_FGC-A+LPS) and incubated with the nanodisc-embedded LptD^I230*pBpa*^E-YifL complex in the presence ATP over a time course. Upon exposure to ultraviolet (UV) radiation, we observed YifL crosslinked to LptD for all three samples. Interestingly, we found that in the presence of ATP and LPS, the ratio of crosslinking decreased with increasing incubation time (Fig. 7A). The results suggest that both LPS and ATP hydrolysis by LptB_2_FG are required for YifL OM translocation. The constant levels of LptD×YifL crosslinks for the nanodisc-embedded LptB_2_FGC-A+YifL sample are consistent with our earlier conclusion that LptB_2_FGC alone can not transfer YifL to LptDE (Fig. 7A).

**Fig. 7.**
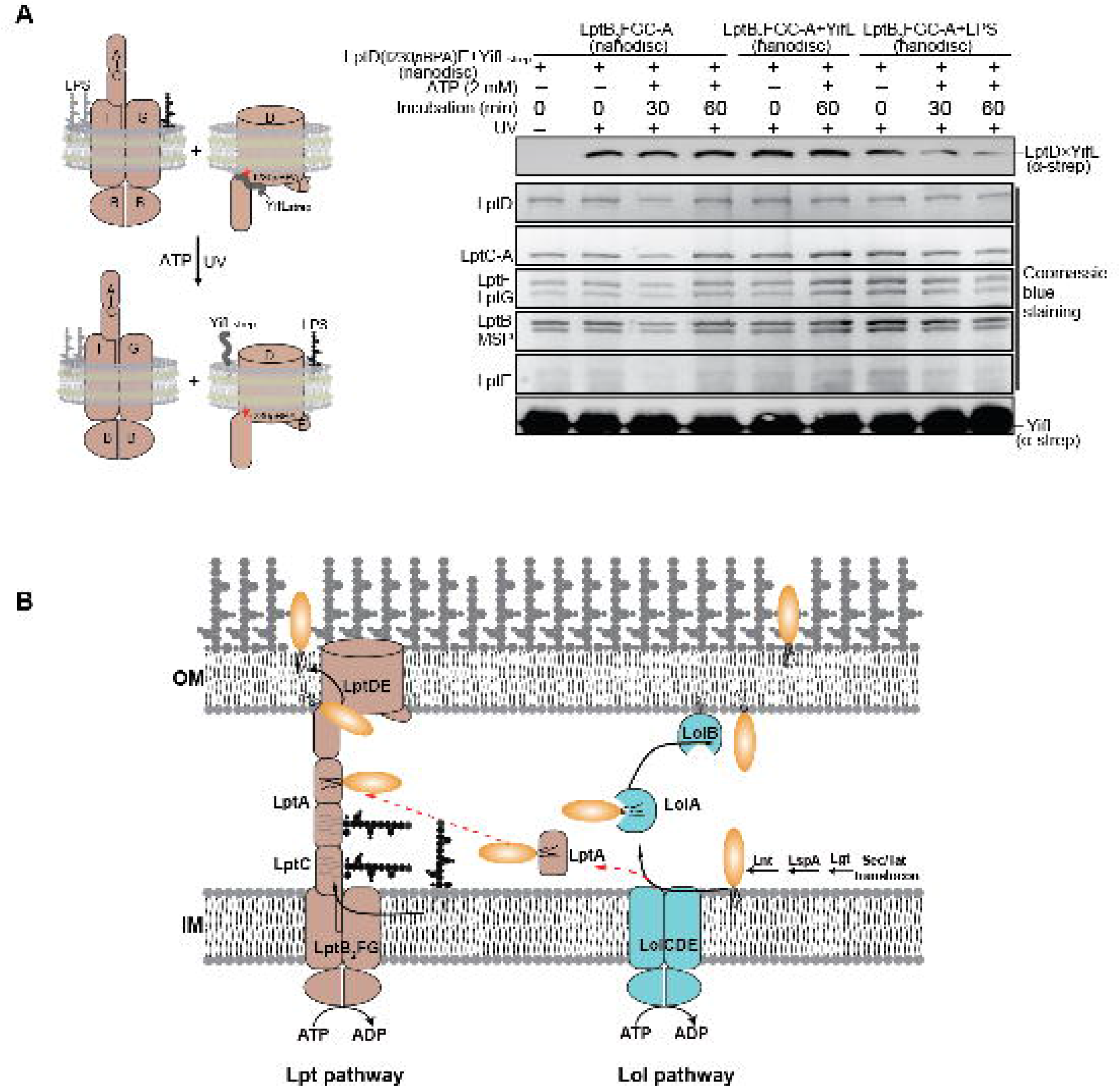
YifL translocates across the OM in an ATP- and LPS-dependent manner. (**A**) *In vitro* LPS membrane-to-membrane transport assay showing that YifL translocates across the OM in an ATP- and LPS-dependent manner. Four different types of nanodiscs were prepared for the assay: LptB_2_FGC-A, LptB_2_FGC-A+YifL, LptB_2_FGC+LPS and LptD^I230*p*Bpa^E-YifL_strep_. The LptD×LPS crosslinks decreased in an ATP- and time-manner only when nanodisc-embedded LptD^I230*p*Bpa^E-YifL_Strep_ was incubated with the nanodiscs containing both LptB_2_FGC and LPS. Cartoons show experimental designs of the *in vitro* LPS transport assay. (**B**) The proposed model of SLP transport to the cell surface. The red dash line indicates that LptA crosstalks to the Lol pathway during OM biogenesis.

## Discussion

In this study, we identified an inventory of *E. coli* lipoproteins that can be transported to the cell surface via LptDE, thereby for the first time we demonstrated that LptDE may function as an OM-transporter for certain lipoproteins. We further show that LptA, a periplasmic component of the Lpt pathway, can shuttle lipoproteins extracted by LolCDE through the periplasm, thus creating a crosstalk between the Lol and Lpt pathways during OM biogenesis. Based on our findings, we propose a previously unidentified sorting route for lipoprotein surface presentation (Fig. 7B). Lipoproteins including SLPs are first extracted from the IM by LolCDE. Both periplasmic chaperones LolA and LptA can accept lipoproteins upon their release from LolCDE. The LolA-bound lipoproteins target to the inner leaflet of the OM via the classical Lol pathway. One the other hand, LptA-bound lipoproteins are incorporated into the Lpt pathway by bridging LptB_2_FGC with LptDE, thereby transported to the cell surface. Third, lipoprotein surface translocation relies on energy provided by ATP hydrolysis (by LptB_2_FG) and LPS to push lipoprotein through to their OM destination^24,42^.

In contrast to a number of previously-reported surface-exposed lipoproteins that require either a dedicated translocon ^44–46^ or the BAM complex ^47,48^ for surface translocation, the SLPs identified here share a same OM translocon, namely LptDE and periplasmic chaperone, LptA. These SLPs, and LPS likewise, all lack large protein domains that can potentially provide energy to drive OM translocation via folding, in an energy-deficient milieu^49–51^. In this regard, it is not surprising that these SLPs utilize the trans-periplasmic scaffold of the Lpt pathway, in which the cytoplasmic ATP serves as energy source driving the SLPs through the hydrophobic channel to the cell surface. However, structure and/or sequence features that dictate surface exposure of a specific lipoprotein await further clarification.

Our *pa*LptDE-YifL structure put forward an alternative mechanism of how a subset of lipoproteins are transported across and anchor to the outer leaflet of the OM. First, the *pa*LptDE-YifL structure snapshots an intermediate state of a lipoprotein en route to the cell surface, demonstrating that the hydrophobic acyl chains and hydrophilic proteinaceous moiety pass through the OM via the exterior and interior of the LptD β-barrel, respectively. Second, the clogged YifL in *pa*LptDE structure suggests a final destination of YifL at the outer leaflet of the OM. This highlights a critical role of the two pairs of conserved interdomain disulfide bonds within LptD in preventing its cargo, LPS or lipoproteins, from mis-localization in the inner leaflet of the OM. Third, the closed conformation of the β-jellyroll domain in the *pa*LptDE-YifL structure, in comparison to the apo- structures, casts doubt on the notion that substrates in the Lpt pathway are transported across the periplasm in a continuous stream, in which scenario the *pa*LptDE-YifL complex would be able to bind multiple substrates simultaneously. Indeed, results from native MS analysis of the *pa*LptDE and *sf*LptDE samples show that LptDE binds lipoproteins at unit stoichiometry, consistent with crystallographic studies. Furthermore, we emphasize that the dynamic nature of the LptD β-jellyroll domains is critical for squeezing substrates by concerted conformational changes of the OM exporter.

LPS and lipoproteins are involved in vital physiological activities of Gram-negative bacteria. Inhibition of either the Lpt or Lol pathways with synthetic chemicals is lethal^52–54^. *pa*LptDE has been demonstrated as an important drug target^52,55^. The structures of *pa*LptDE-YifL and apo-*pa*LptDE we present here can therefore provide structural guidance for optimization of the known peptidomimetic antibiotics against drug-resistant *P. aeruginosa.* In addition, it can also aid in drug development targeting surface-exposed lipoproteins. Additionally, the SLPs we identified in this study expand our toolkit of surface display strategies as fusion partners for protein of interest.

## Methods and Materials

### Protein expression, purification and crystallization

We cloned *lptDE* genes from two different species into vector pBAD22 under control of the arabinose promoter. The plasmids contained the genes for *lptD* and *lptE* in this order, each with a separate translation-initiation site. The full-length *lptD* and *lptE* genes were amplified from *Pseudomonas aeruginosa* PAO1-LAC (47085D-5) and *Shigella flexneri* (700030D) genomic DNAs (ATCC) by polymerase chain reactions (PCR). All the generated plasmids (pBAD22- *palptDE* and pBAD22-*sflptDE*) contained a hexahistidine tag at the C-terminus of LptE that was introduced in the PCR primer to facilitate subsequent affinity purification. The plasmid pBAD22-*lptDE* was then transformed into the protease-deficient *E. coli* strain SF100 [KS272Δ*(ompT-entF)]* (ATCC) for co-expression. Protein expression was induced by the addition of 0.5% L-arabinose for 12 hrs at 26°C when the O.D._600_ of the culture reached about 1.0. Cells were harvested by centrifugation at 4,500 g for 30 min at 4°C. Cell pellets were resuspended in 1×PBS (pH 7.4), lysed by a single passage through a French Press (JN-3000 PLUS, China) at 16,000 psi, and centrifuged at 39,000 g for 45 min at 4°C to collect the total cell membranes. The total membranes were solubilized with a buffer containing 1×PBS (pH 7.4) and 0.5% *N*-lauroylsarcosine sodium (Sigma-Aldrich) for 1 hr at 4°C. The outer membranes were isolated by centrifugation at 39,000 g for 1 hr at 4°C and were further solubilized for 1 hr with buffer A [20 mM Tris-HCl (pH 8.0), 150 mM NaCl, 20 mM imidazole and 1% *N,N*-Dimethyldodecylamine *N*-oxide (LDAO)]. The supernatant was collected after centrifugation at 39,000 g for 1 hr at 4°C and incubated with pre-equilibrated Ni-NTA agarose beads for 2 hrs at 4 °C. Protein-bound Ni-NTA agarose beads were rinsed with buffer B [20 mM Tris-HCl (pH 8.0), 150 mM NaCl, 30 mM imidazole and 0.2% LDAO], and detergent exchange was performed with buffer C [20 mM Tris-HCl (pH 8.0), 150 mM NaCl, and 1% tetraethylene glycol monooctyl ether (C_8_E_4_)]. The LptDE complex was eluted from the Ni-NTA agarose beads using buffer D [20 mM Tris-HCl (pH 8.0), 150 mM NaCl, 200 mM imidazole and 1% C_8_E_4_]. The eluted LptDE complex was subsequently applied to a Resource-Q column (GE Healthcare), and followed by a Superdex^™^ 200 10/300 size exclusion column (GE Healthcare) that was pre-equilibrated with 20 mM Tris-HCl (pH 8.0), 150 mM NaCl and 0.6% C_8_E_4_. The purified *pa*LptDE and *sf*LptDE samples were shown to contain lipoprotein YifL as revealed by 15% SDS-PAGE analysis. Normally, we obtained approximately 1.5 mg protein for *pa*LptDE complex from about 80 liters of Terrific Broth (TB) culture, which is a 10-fold lower than the yield of *sf*LptDE.

Crystallization was conducted at 16°C using the hanging drop vapor diffusion method, mixing 1 μl each of *pa*LptDE-YifL (15 mgml^-1^) and reservoir solutions at a ratio of 1:1. Initial crystallization condition was found using a broad screening. After optimization, the best crystals were obtained in a buffer containing 175 mM citric acid/Bis-Tris (pH 5.0), 17% (v/v) PEG3350 and 3% methanol (as additive). The *pa*LptDE-YifL crystals appeared overnight and grew to their final size approximately 100×120×60 μm in about one week. They belong to space group P2_1_ and the best crystals diffracted to 3.0 Å at synchrotron. The apo-*pa*LptDE protein was purified from the *yifL*-depleted *E. coli* SF100 strain using a similar protocol as described for *pa*LptDE-YifL. The apo-*pa*LptDE crystals were obtained in a condition that contained 2% tacsimate (pH5.0), 16% PEG3350 and 100 mM sodium citrate tribasic dehydrate at 16°C and the best crystals diffracted to 3.3 Å. The crystals belong to space group P2_1_2_1_2_1_ with two apo-*pa*LptDE complexes in one asymmetric unit. All the crystals were flash frozen in liquid nitrogen by addition of 20% glycerol into the reservoir solutions for data collection.

### Structure determination and refinement

All diffraction data were collected at Shanghai Synchrotron Radiation Facility (SSRF, Shanghai, China). X-ray data were processed with HKL2000^56^. For both *pa*LptDE-YifL and *pa*LptDE, an initial molecular replacement solution was found using Phaser with the jellyroll-truncated *pa*LptDE (PDB code: 5IVA) as search model ^57^. The electron densities for the jellyroll domains of *pa*LptD were resolved after molecular replacement, followed by several rounds of manual building for the jellyroll domain of *pa*LptD with Coot ^58 57^β-barrel lumen exhibited polypeptide features with discernible bulky side chains, which allowed us to manually build YifL into the structure. Further SDS-PAGE analysis using dissolved crystals confirmed the presence of YifL in the crystals. Refinements of the *pa*LptDE-YifL and apo-*pa*LptDE datasets were performed using Phenix and CCP4 refmac5 ^59^. *Pa*LptDE-YifL and *pa*LptDE structures were refined to 3.0-Å and 3.3-Å resolution with R_work_/R_free_=0.22/0.26 and R_work_/R_free_=0.22/0.26, respectively. The poorly defined residues (matured proteins) in the best-refined structures of *pa*LptDE-YifL (*pa*LptD: 1-81; YifL: 12-46) and apo-*pa*LptDE (*pa*LptD: 1-70) presumably indicate conformational flexibility and partial degradation of the *pa*LptD N-terminus. All structure figures were rendered using PyMOL ^60^. The detailed refinement statistics for both *pa*LptDE-YifL and *pa*LptDE structures are listed in Table 1.

### Construction of the *yifL*-depleted *E. coli* SF100 strain

To delete the *yifL* gene in *E. coli* sf 100 strain, the linear fragments were amplified by PCR using primers P1 (5’-TCCTGCGATG ATAGAAAGCA GAAAGCGATG AACTTTACAG GCAATCCATA GTGTAGGCTGGAGCTGCTTC-3’) and P2 (5’-TTCGAGAACT GCATCATTTA CTCCAATCAC GCGGGTACAG AAACTGACTT CATATGAATATCCTCCTTAG-3’) with 50-nt extensions that are homologous to regions adjacent to the *yifL* gene using the plasmid pKD3 as template. PCR fragments were then transformed by electroporation into the *E. coli* SF100 strain that carries the pKD46 plasmid to obtain *yifL*::Cm. Mutants were colony-purified once non-selectively at 37 °C to eliminate pKD46. The Cm cassette was then removed by introducing the helper plasmid pCP20 into the mutant *yifL*::Cm and colony-purified once non-selectively at 43 °C. The *yifL*-depleted *E. coli* SF100 strain was used for over-expressing *pa*LptDE proteins to obtain apo-*pa*LptDE crystals.

### Native mass spectrometry

Prior to mass spectrometry (MS) analysis, the purified *pa*LptDE or *sf*LptDE protein samples were buffer exchanged into 150 mM ammonium acetate (pH 8.0) containing 0.5% C_8_E_4_ (v/v) on a Superdex 200 gel filtration column (GE Healthcare). LptDE samples were subsequently introduced via gold-coated nanospray capillaries prepared in house and MS spectra were recorded on a Synapt G2-Si instrument (Waters, Manchester, UK) modified for high mass detection. Optimized instrument parameters include capillary voltage 1.8 kV, sampling cone 150 V, sampling offset 100 V, trap collision energy 150 V, transfer collision energy 35 V, trap gas flow 5.0 mlmin^-1^. MS data were analyzed using MassLynx version 4.1 (Waters, Manchester, UK).

### Pulldown and immunoblotting assay

To confirm the interactions between *sf*LptDE and the nine identified lipoproteins (and lipoprotein BamE as negative control), plasmids contained the genes for *sflptD, sflptE_6×His_* and a lipoprotein with a C-terminal strep tag II in this order, each with a separate translation-initiation site, were cloned in the pBAD22 vector. The constructed plasmids were individually transformed into *E. coli* SF100 cells for co-expression. Protein expression and purification followed a similar procedure as described for *pa*LptDE, but only 1 liter of LB culture was prepared for each construct. All the cell lysates were prepared from the total membranes solubilized with 1% LDAO. Each of the purified *sf*LptDE_6×His_ proteins and its cell lysate were subjected to SDS-PAGE (15%) analysis.

After electrophoresis, the proteins were transferred to a PVDF membrane and blocked using TBST buffer (20 mM Tris-HCl, pH 8.0, 150mM NaCl, 0.05% Tween-20) that contained 5% skim milk for 1 hr. The PVDF membrane was then incubated with anti-Strep tag II mouse monoclonal antibody (1:3500) (Beijing Xuheyuan Biotech) at room temperature for 1 hr and subsequently washed with TBST buffer twice and further incubated with anti-mouse IgG (H+L)-HRP (1:3500) (LabLead) at room temperature for 1 hr. PVDF membranes were exposed using enhanced chemiluminescence reagents (EasySee Western Blot Kit, Trans^™^).

### Immunofluorescence assay

To carry out immunofluorescence assay, each of the identified lipoproteins and BamB (as negative control) with a C-terminal His tag were cloned into the pET22b vector and was then transformed into *E. coli* SF100 cells. For each sample, 1 ml *E. coli* cell culture (O.D._600_=1.0) was loaded into a 1.5-ml Eppendorf tube. After removing the supernatant by centrifugation, each cell pellet was washed twice with PBS. The cell pellet for each sample was then resuspended with 1 ml PBS, and 200-μl suspension for each sample was transferred into the glass bottom cell culture dish (NEST, ø15 mm) that was pretreated with 0.01% poly-L-lysine. Following incubation for 5 min, the redundant suspension was discarded. The bacteria attached to the dish were fixed by addition of 200 μl 4% (w/v) paraformaldehyde in PBS for 2 hrs, followed by wash twice with 1 ml PBS for each sample.

The fixed bacteria were then blocked in 10% normal goat serum in PBS for 30 min. After being blocked, the samples were incubated with primary antibody (mouse anti-His, 1:100, CMCTAG) for 2 hrs, and then washed with PBS three times, 5 min each. The samples were then stained with Alexa Fluor 546-conjugated secondary antibody (goat anti-mouse, 1:200, Invitrogen) for 1 hr. After being washed for three times with PBS, they were mounted in Moviol Mounting Media with 4’,6’-diamidino-2-phenylindole (DAPI, blue-fluorescing), a membrane-penetrating fluorescent dye that binds strongly to adenine-thymine rich regions in DNA. For *E coli* cells that harbored the pET22b-*bamB* plasmid, an addition step was performed by treating with 0.5% Triton X-100 and 5 mM EDTA for 30 min to punch the membrane before proceeding to block with 10% normal goat serum in PBS.

The bacteria images were acquired on the Delta Vision OMX V3 image system (GE Healthcare) with a ×100/1.40 NA oil-immersion objective lens (Olympus UPlanSApo) and a camera (Evolve 512×512, Photometrics). Images were processed and analyzed using Image J software (NIH). The experiments were performed in duplicate and repeated three times, and the results are representative of replicates.

### Dot blot assay

Dot blot assays were carried out similarly as described in a previous study ^61^. Briefly, *E. coli* SF100 cells that carried either YifL or BamB (as negative control)-expressing plasmid (pET22b-*yifL*_6×His_ or pET22b-*bamB*_6×His_) were grown to OD_600_ = 1.0. 1 ml of cell culture was withdrawn for each sample, and was washed twice with PBS (10 mM Na_2_HPO_4_, 1.8 mM KH_2_PO_4_, 2.7 mM KCl, 137 mM NaCl, pH 7.4). Cell lysates were prepared by sonication of the cell suspension in PBS supplemented with 10 mM EDTA. Three microliters of cell suspension (whole cell) or cell lysate were spotted on an Immobilon-P^SQ^ transfer membrane (Merck Millipore Ltd.) and air dried. Membranes were blocked with 1% (w/v) skim milk in PBS for 30 min at room temperature and probed with anti-His mouse monoclonal antibody (TansGen Biotech) for 1 hr at room temperature. The membranes were washed three times for 5 min with PBS and probed with goat anti-mouse IgG (H+L) HRP-conjugated secondary antibody (1:3500) (LabLead) for 1 hr at room temperature. Membranes were exposed using enhanced chemiluminescence reagents (EasySee Western Blot Kit, Trans^™^). The experiments were performed in triplicate and repeated at least three times, and the results are representative of replicates.

### Molecular dynamics (MD) simulation

To carry out MD simulations, we first generated two *pa*LptDE complex models (apo-*pa*LptDE and *pa*LptDE-YifL) based on the apo-*pa*LptDE and *pa*LptDE-YifL crystal structures. The structure of the missing residues in the *pa*LptDE-YifL model was built based on the apo-*pa*LptD crystal structure. The two pairs of nonconsecutive disulfide bonds of LptD such as Cys107-Cys831 and Cys243-Cys832 were built for each model, and the first Cys residue of lipoproteins *pa*LptE and YifL was triacylated. Each of the apo-*pa*LptDE and *pa*LptDE-YifL complex models was then embedded into an *E.coli* K12 OM comprised of K12 LPS in the outer leaflet and an inner leaflet phospholipid mixture of 1-palmitoyl (16:0)-2-palmitoleoyl (16:1 cis-9)-phosphatidylethanolamine, 1-palmitoyl (16:0)-2-vacenoyl (18:1 cis-11)-phosphatidylglycerol, and 1,10-palmitoyl-2,20-vacenoyl cardiolipin at a ratio of 15:4:1 An assembly of all components was made by a systematic protocol of CHARMM-GUI Membrane Builder (Kumar et al., 2007).^62^ Neutralizing Ca^2+^ ions were initially placed in the LPS region to mediate crosslinking networks with negatively charged groups, and 150 mM KCl ions were added with TIP3P water molecules ^64^.

For each system, the preliminary NAMD simulations with the CHARMM 36 force field for proteins^66^, carbohydrates^67^, lipids^68^ and LPS^69^ were performed over 20 ns for the preparation for Anton simulations. For NAMD simulations, each system was equilibrated with 75-ps NVT (constant particle number, volume, and temperature) dynamics with 1-fs time step, followed by 300-ps NPT (constant particle number, pressure, and temperature) dynamics with a 2-fs time step and gradually relaxed positional (protein) planar (membrane), and dihedral (carbohydrate) harmonic restraints. We then performed additional 20-ns NPT dynamics without any restraint except the dihedral restraint. For the data analysis presented in this paper, 1-μs Anton simulations^70^ were performed with *Multigrator* integrator^71^ resulting in a framework for building constant temperature and pressure integrators. The temperature and the pressure for all simulations in this study were held at 303.15 K and 1 bar, respectively.

### Nanodisc preparation

*E. coli* Total Lipid Extract (Avanti Lipids) was solubilized in chloroform, dried under nitrogen to form a thin lipid film. The lipid film was hydrated and resuspended at a concentration of 25 mM Total Lipid in 250 mM sodium cholate. Each membrane protein complex (LptDE, LptDE-YifL, LptB_2_FGC, LptB_2_FGC-A or LolCDE), MSP1D1 membrane scaffold protein ^72^, and Total Lipid were mixed at a molar ratio of 1:2.4:80 in a buffer containing 15 mM sodium cholate and incubated for 30 min at 4 °C. Detergents were removed by incubation with 0.8 mg ml^-1^ Bio-Beads SM2 (Bio-Rad) overnight at 4 °C. Nanodisc-embedded complexes were further purified using a Superose6 increase 10/300GL column (GE Healthcare) in a buffer containing 20 mM Tris-HCl (pH 8.0) and 150 mM NaCl. Peak fractions were combined and concentrated to approximately 0.6 mg ml^-1^.

### *In vivo* photocrosslinking

To test whether YifL cross-links to LptDE in *E. coli,* expression plasmids pBAD22-*lptDE* and pET28a-*yifL*_Strep_ were constructed. pBAD22-*lptDE* plasmids contain an amber (TAG) codon at either LptD^I230^ or LptD^Y112^ for incorporation of *p*Bpa. *E. coli* BL21 (DE3) cells were transformed with three plasmids pSup-BpaRS-6TRN22, pBAD22-*lptDE* and pET28a-*yifL*_Strep_ simultaneously for protein expression. The transformed *E. coli* cells were grown at 37 °C in LB in the dark, supplemented with ampicillin sodium (30 μg ml^-1^), kanamycin (15 μg ml^-1^) and chloramphenicol (15 μgml^-1^). When the culture reached O.D._600_ = 1.0, *p*Bpa was added into LB at a final concentration of 0.5 mM. After 1 h, protein expression was induced by the addition of 0.05 mM IPTG and 0.05% L-arabinose at 18 °C for 4 hrs. 200 μl culture aliqouts withdrawn were used either directly or exposed to UV light (365 nm, 100 W; Thermo Fisher Scientific) for 10 min. Proteins were separated on a 15% SDS-PAGE gel, and crosslinks were detected with Strep tag II monclonal antibody (Thermo Fisher Scientific).

To investigate whether LptA cross-links to YifL, LolB or NlpA, expression plasmids pQlinkN-*lptA*, pET28a-*yifL*_Strep_, pET28a-*lolpB*_Strep_ and pET28a-*nlpA*_Strep_, were constructed. pQlinkN-*lptA* plasmids contain an amber (TAG) codon at either LptA^F95^ or LptA^I36^ for incorporation of *p*Bpa. *E. coli* BL21 (DE3) cells were transformed with three plasmids pSup-BpaRS-6TRN22, pQlinkN-*lptA* and pET28a-*yifL*_Strep_ (pET28a-*lolB*_Strep_ or pET28a-*nlpA*_Strep_). The transformed *E. coli* cells were grown at 37 °C in the dark, supplemented with ampicillin sodium (30 μg ml^-1^), kanamycin (15 μg ml^-1^) and chloramphenicol (15 μg ml^-1^). When the cultures reached O.D._600_ = 1.0, *p*Bpa was added to LB at a final concentration of 0.5 mM. After 1 hr, protein expression was induced by the addition of 0.05 mM IPTG at 18 °C for 4 hrs. Photocrosslinks were detected in a similar way as described above.

### *In vitro* photocrosslinking assay

Expression and purification of YifL for nanodisc reconstitution. The *E. coli yifl* gene (along with the C-terminal Strep tag II coding sequence) was cloned into vector pET28a. The generated pET28a-*yifL*_Strep_ plasmid was transformed into *E. coli* BL21 (DE3) strain. The pET28a-YifL_Strep_ - harboring *E. coli* cells were grown at 37 °C in LB with 35 μgml^-1^ kanamycin. Protein expression was induced by addition of 0.1 mM IPTG at 25 °C for 12 hrs when the optical density of the culture reached 1.0 at 600 nm. YifL was purified from the membrane fractions by using 1.0% DDM. After the detergent-solubilized supernatants were incubated with pre-equilibrated strep-tactin beads for 1 hr, YifL_Strep_ protein was eluted with wash buffer containing 2.5 mM d-Desthiobiotin (Sigma).

YifL(or LPS), *p*Bpa-containing LptB_2_FGC proteins (or *p*Bpa-containing LolCDE proteins), MSP1D1 and Total Lipid were mixed at a molar ratio of 1:1:2.4:80 and incubated for 30 min at 4 °C. Detergents were removed by incubation with 0.8 mg ml^-1^ Bio-Beads SM2 (Bio-Rad) overnight at 4 °C. Nanodisc-embedded complexes were further purified using a Superose6 increase 10/300GL column (GE Healthcare) in a buffer containing 20 mM Tris-HCl (pH 8.0) and 150 mM NaCl. Peak fractions were combined and concentrated to approximately 0.6 mg ml^-1^. The reconstituted nanodisc samples were incubated with or without 2 mM ATP (and 2 mM MgCl_2_) at room temperature for various lengths of time (0-10 min), followed by UV radiation for 10 min when needed.

To carry out YifL transfer assay, nanodisc-embedded LolCDE-YifL complexes were mixed with either LolA(W70pBpa) or LptA(F95pBpa) at a molar ratio of 1:1 at room temperature. After addition of 2 mM ATP (and 2 mM MgCl_2_), the mixture was incubated for 10 min at room temperature, followed by UV radiation for 10 min when needed. LPS/YifL membrane-to-membrane transport was performed using a similar procedure but using nanodiscs that contained different membrane protein complexes.

For western blot detection of crosslinks, the reaction mixtures were first separated by 15% SDS–PAGE gel, and transferred to PVDF membranes (Bio-Rad). Following blocking for 1 hr with TBST buffer [20 mM Tris-HCl (pH 8.0), 150mM NaCl, 0.1% Tween-20] containing 8% skim milk, the PVDF membrane was incubated with either Strep tag II antibody (1:3000 dilution, AP1013a, YTHX, China) or anti-*E. coli* LPS antibody (1:3000 dilution, ab35654, Abcam) at room temperature for 1 hr. After washing with TBST buffer three times, the PVDF membrane was incubated with goat anti-rabbit or goat anti-mouse horseradish per-oxidase (HRP)-conjugated secondary antibody (1:5000 dilution, LABLEAD, China) at room temperature for 1 hr. YifL or LPS crosslinks were visualized using an enhanced chemiluminescence detection kit (Applygen, China).

## Author contributions

Y.H. conceived and supervised the project. Q.L., C.W., S.Q., S.Y., L.C., S.K., and K.W. performed the experiments. C.W., S.Q., Q.L., J.Z. and Y.H. collected diffraction data, built the model and refined the structure; K.W. and M.Z. performed the native mass spectrometry analysis; L.C. and L. M. carried out the immunofluorescence assays; S.K. and W.I. contributed to the MD simulation studies; F.W., X.L., and M.H. provided materials. Q.L. M.Z. and Y.H. wrote the manuscript with the input from all authors. The coordinates and diffraction data of apo-*pa*LptDE and *pa*LptDE-YifL crystal structures have been deposited in the Protein Data Bank with the accession codes 8H1S and 8H1R, respectively. The authors declare no competing financial interests.

## Acknowledgments

The authors would like to thank Dr. Chérine Bechara at University of Oxford_for preliminary native mass spectrometry analysis of the *pa*LptDE protein sample; Dr. Marcin Grabowicz at Emory University School of Medicine provided the *lolA*-depleted MG3324 (NR754 *Δlpp ΔrcsB zii::*Tn10 *cpxA24 ΔlolA*) *E. coli* strain. Ms. Shuoguo Li from Center for Biological Imaging (CBI), Institute of Biophysics, Chinese Academy of Sciences (CAS), for help on taking and analyzing SIM images. The diffraction data were collected at beamlines BL19U and BL17U of the Shanghai Synchrotron Radiation Facility (SSRF, China). This work was supported by grants from the Strategic Priority Research Program of the Chinese Academy of Sciences (XDB37020201 to Y. H.), the Chinese Academy of Sciences (CAS201973), the National Natural Science Foundation of China (31625009 to Y.H.), the China Postdoctoral Science Foundation (BX20190356 to Q.L.) and the National Science Foundation (MCB-1727508 and MCB-1810695 to W.I.).

## References

1 Nikaido, H. Molecular basis of bacterial outer membrane permeability revisited. Microbiology and molecular biology reviews: MMBR 67, 593–656 (2003).

2 Konovalova, A., Kahne, D. E. & Silhavy, T. J. Outer Membrane Biogenesis. Annual review of microbiology 71, 539–556, doi:10.1146/annurev-micro-090816-093754 (2017).

3 May, K. L. & Silhavy, T. J. Making a membrane on the other side of the wall. Biochimica et biophysica acta 1862, 1386–1393, doi:10.1016/j.bbalip.2016.10.004 (2017).

4 Bos, M. P., Robert, V. & Tommassen, J. Biogenesis of the gram-negative bacterial outer membrane. Annual review of microbiology 61, 191–214, doi:10.1146/annurev.micro.61.080706.093245 (2007).

5 Chng, S. S., Ruiz, N., Chimalakonda, G., Silhavy, T. J. & Kahne, D. Characterization of the two-protein complex in Escherichia coli responsible for lipopolysaccharide assembly at the outer membrane. Proceedings of the National Academy of Sciences of the United States of America 107, 5363–5368, doi:10.1073/pnas.0912872107 (2010).

6 Aliprantis, A. O. et al. Cell activation and apoptosis by bacterial lipoproteins through toll-like receptor-2. Science 285, 736–739 (1999).

7 Vijay, K. Toll-like receptors in immunity and inflammatory diseases: Past, present, and future. International immunopharmacology 59, 391–412, doi:10.1016/j.intimp.2018.03.002 (2018).

8 Poltorak, A. et al. Defective LPS signaling in C3H/HeJ and C57BL/10ScCr mice: mutations in Tlr4 gene. Science 282, 2085–2088 (1998).

9 Sperandeo, P. et al. Functional analysis of the protein machinery required for transport of lipopolysaccharide to the outer membrane of Escherichia coli. Journal of bacteriology 190, 4460–4469, doi:10.1128/JB.00270-08 (2008).

10 Okuda, S., Sherman, D. J., Silhavy, T. J., Ruiz, N. & Kahne, D. Lipopolysaccharide transport and assembly at the outer membrane: the PEZ model. Nature reviews. Microbiology 14, 337–345, doi:10.1038/nrmicro.2016.25 (2016).

11 Sperandeo, P. et al. Characterization of lptA and lptB, two essential genes implicated in lipopolysaccharide transport to the outer membrane of Escherichia coli. Journal of bacteriology 189, 244–253, doi:10.1128/JB.01126-06 (2007).

12 Ruiz, N., Gronenberg, L. S., Kahne, D. & Silhavy, T. J. Identification of two inner-membrane proteins required for the transport of lipopolysaccharide to the outer membrane of Escherichia coli. Proceedings of the National Academy of Sciences of the United States of America 105, 5537–5542, doi:10.1073/pnas.0801196105 (2008).

13 Sherman, D. J. et al. Decoupling catalytic activity from biological function of the ATPase that powers lipopolysaccharide transport. Proceedings of the National Academy of Sciences of the United States of America 111, 4982–4987, doi:10.1073/pnas.1323516111 (2014).

14 Tran, A. X., Dong, C. & Whitfield, C. Structure and functional analysis of LptC, a conserved membrane protein involved in the lipopolysaccharide export pathway in Escherichia coli. The Journal of biological chemistry 285, 33529–33539, doi:10.1074/jbc.M110.144709 (2010).

15 Luo, Q. et al. Structural basis for lipopolysaccharide extraction by ABC transporter LptB2FG. Nature structural & molecular biology 24, 469–474, doi:10.1038/nsmb.3399 (2017).

16 Dong, H., Zhang, Z., Tang, X., Paterson, N. G. & Dong, C. Structural and functional insights into the lipopolysaccharide ABC transporter LptB2FG. Nature communications 8, 222, doi:10.1038/s41467-017-00273-5 (2017).

17 Li, Y., Orlando, B. J. & Liao, M. Structural basis of lipopolysaccharide extraction by the LptB2FGC complex. Nature 567, 486–490, doi:10.1038/s41586-019-1025-6 (2019).

18 Owens, T. W. et al. Structural basis of unidirectional export of lipopolysaccharide to the cell surface. Nature 567, 550–553, doi:10.1038/s41586-019-1039-0 (2019).

19 Tang, X. et al. Cryo-EM structures of lipopolysaccharide transporter LptB2FGC in lipopolysaccharide or AMP-PNP-bound states reveal its transport mechanism. Nature communications 10, 4175, doi:10.1038/s41467-019-11977-1 (2019).

20 Dong, H. et al. Structural basis for outer membrane lipopolysaccharide insertion. Nature 511, 52–56, doi:10.1038/nature13464 (2014).

21 Qiao, S., Luo, Q., Zhao, Y., Zhang, X. C. & Huang, Y. Structural basis for lipopolysaccharide insertion in the bacterial outer membrane. Nature 511, 108–111, doi:10.1038/nature13484 (2014).

22 Botos, I. et al. Structural and Functional Characterization of the LPS Transporter LptDE from Gram-Negative Pathogens. Structure 24, 965–976, doi:10.1016/j.str.2016.03.026 (2016).

23 Sherman, D. J. et al. Lipopolysaccharide is transported to the cell surface by a membrane-to-membrane protein bridge. Science 359, 798–801, doi:10.1126/science.aar1886 (2018).

24 Okuda, S., Freinkman, E. & Kahne, D. Cytoplasmic ATP hydrolysis powers transport of lipopolysaccharide across the periplasm in E. coli. Science 338, 1214–1217, doi:10.1126/science.1228984 (2012).

25 Okuda, S. & Tokuda, H. Lipoprotein sorting in bacteria. Annual review of microbiology 65, 239–259, doi: 10.1146/annurev-micro-090110-102859 (2011).

26 Konovalova, A. & Silhavy, T. J. Outer membrane lipoprotein biogenesis: Lol is not the end. Philosophical transactions of the Royal Society of London. Series B, Biological sciences 370, doi:10.1098/rstb.2015.0030 (2015).

27 Narita, S. I. & Tokuda, H. Bacterial lipoproteins; biogenesis, sorting and quality control. Biochimica et biophysica acta 1862, 1414–1423, doi:10.1016/j.bbalip.2016.11.009 (2017).

28 Wilson, M. M. & Bernstein, H. D. Surface-Exposed Lipoproteins: An Emerging Secretion Phenomenon in Gram-Negative Bacteria. Trends in microbiology 24, 198–208, doi:10.1016/j.tim.2015.11.006 (2016).

29 Cowles, C. E., Li, Y., Semmelhack, M. F., Cristea, I. M. & Silhavy, T. J. The free and bound forms of Lpp occupy distinct subcellular locations in Escherichia coli. Molecular microbiology 79, 1168–1181, doi:10.1111/j.1365-2958.2011.07539.x (2011).

30 Sukupolvi, S. & O’Connor, C. D. TraT lipoprotein, a plasmid-specified mediator of interactions between gram-negative bacteria and their environment. Microbiological reviews 54, 331–341 (1990).

31 Grabowicz, M. Lipoprotein Transport: Greasing the Machines of Outer Membrane Biogenesis: Re Examining Lipoprotein Transport Mechanisms Among Diverse Gram-Negative Bacteria While Exploring New Discoveries and Questions. Bioessays 40, 1700187 (2018).

32 Braun, V. & Bosch, V. Sequence of the murein-lipoprotein and the attachment site of the lipid. European journal of biochemistry 28, 51–69 (1972).

33 Botte, M. et al. Cryo-EM structures of a LptDE transporter in complex with Pro-macrobodies offer insight into lipopolysaccharide translocation. Nature communications 13, 1–10 (2022).

34 Moehle, K. et al. Solution Structure and Dynamics of LptE from Pseudomonas aeruginosa. Biochemistry 55, 2936–2943, doi:10.1021/acs.biochem.6b00313 (2016).

35 Chng, S. S. et al. Disulfide rearrangement triggered by translocon assembly controls lipopolysaccharide export. Science 337, 1665–1668, doi:10.1126/science.1227215 (2012).

36 Han, L. et al. Structure of the BAM complex and its implications for biogenesis of outer-membrane proteins. Nature structural & molecular biology 23, 192–196, doi:10.1038/nsmb.3181 (2016).

37 Wu, T. et al. Identification of a multicomponent complex required for outer membrane biogenesis in Escherichia coli. Cell 121, 235–245, doi:10.1016/j.cell.2005.02.015 (2005).

38 Li, X., Gu, Y., Dong, H., Wang, W. & Dong, C. Trapped lipopolysaccharide and LptD intermediates reveal lipopolysaccharide translocation steps across the Escherichia coli outer membrane. Scientific reports 5, 11883, doi:10.1038/srep11883 (2015).

39 Ryu, Y. & Schultz, P. G. Efficient incorporation of unnatural amino acids into proteins in Escherichia coli. Nature methods 3, 263–265, doi:10.1038/nmeth864 (2006).

40 Sharma, S. et al. Mechanism of LolCDE as a molecular extruder of bacterial triacylated lipoproteins. Nature communications 12, 1–11 (2021).

41 Tang, X. et al. Structural basis for bacterial lipoprotein relocation by the transporter LolCDE. Nature structural & molecular biology 28, 347–355 (2021).

42 Grabowicz, M. & Silhavy, T. J. Redefining the essential trafficking pathway for outer membrane lipoproteins. Proceedings of the National Academy of Sciences of the United States of America 114, 4769–4774, doi:10.1073/pnas.1702248114 (2017).

43 Kaplan, E., Greene, N. P., Crow, A. & Koronakis, V. Insights into bacterial lipoprotein trafficking from a structure of LolA bound to the LolC periplasmic domain. Proceedings of the National Academy of Sciences 115, E7389–E7397 (2018).

44 Hooda, Y. et al. Slam is an outer membrane protein that is required for the surface display of lipidated virulence factors in Neisseria. Nature microbiology 1, 16009, doi:10.1038/nmicrobiol.2016.9 (2016).

45 Hooda, Y. & Moraes, T. F. Translocation of lipoproteins to the surface of gram negative bacteria. Current opinion in structural biology 51, 73–79, doi:10.1016/j.sbi.2018.03.006 (2018).

46 Hooda, Y., Shin, H. E., Bateman, T. J. & Moraes, T. F. Neisserial surface lipoproteins: structure, function and biogenesis. Pathogens and disease 75, doi:10.1093/femspd/ftx010 (2017).

47 Konovalova, A., Perlman, D. H., Cowles, C. E. & Silhavy, T. J. Transmembrane domain of surface-exposed outer membrane lipoprotein RcsF is threaded through the lumen of β-barrel proteins. Proceedings of the National Academy of Sciences 111, E4350–E4358 (2014).

48 Oomen, C. J. et al. Structure of the translocator domain of a bacterial autotransporter. The EMBO journal 23, 1257–1266 (2004).

49 Dong, C. et al. Wza the translocon for E. coli capsular polysaccharides defines a new class of membrane protein. Nature 444, 226–229 (2006).

50 Goyal, P. et al. Structural and mechanistic insights into the bacterial amyloid secretion channel CsgG. Nature 516, 250–253 (2014).

51 Cao, B. et al. Structure of the nonameric bacterial amyloid secretion channel. Proceedings of the National Academy of Sciences of the United States of America 111, E5439–5444, doi:10.1073/pnas.1411942111 (2014).

52 Srinivas, N. et al. Peptidomimetic antibiotics target outer-membrane biogenesis in Pseudomonas aeruginosa. Science 327, 1010–1013, doi:10.1126/science.1182749 (2010).

53 Nickerson, N. N. et al. A Novel Inhibitor of the LolCDE ABC Transporter Essential for Lipoprotein Trafficking in Gram-Negative Bacteria. Antimicrob Agents Chemother 62, doi:10.1128/AAC.02151-17 (2018).

54 Narita, S., Tanaka, K., Matsuyama, S. & Tokuda, H. Disruption of lolCDE, encoding an ATP-binding cassette transporter, is lethal for Escherichia coli and prevents release of lipoproteins from the inner membrane. Journal of bacteriology 184, 1417–1422 (2002).

55 Andolina, G. et al. A peptidomimetic antibiotic interacts with the periplasmic domain of LptD from Pseudomonas aeruginosa. ACS chemical biology 13, 666–675 (2018).

56 Otwinowski, Z. & Minor, W. Processing of X-ray diffraction data collected in oscillation mode. Method Enzymol 276, 307–326, doi:Doi 10.1016/S0076-6879(97)76066-X (1997).

57 Adams, P. D. et al. PHENIX: building new software for automated crystallographic structure determination. Acta Crystallogr D 58, 1948–1954, doi:10.1107/S0907444902016657 (2002).

58 Emsley, P. & Cowtan, K. Coot: model-building tools for molecular graphics. Acta Crystallogr D 60, 2126–2132, doi:10.1107/S0907444904019158 (2004).

59 Bailey, S. The Ccp4 Suite - Programs for Protein Crystallography. Acta Crystallogr D 50, 760–763 (1994).

60 DeLano, W. L. PyMOL molecular viewer: Updates and refinements. Abstr Pap Am Chem S 238 (2009).

61 Konovalova, A., Perlman, D. H., Cowles, C. E. & Silhavy, T. J. Transmembrane domain of surface-exposed outer membrane lipoprotein RcsF is threaded through the lumen of beta-barrel proteins. Proceedings of the National Academy of Sciences of the United States of America 111, E4350–4358, doi:10.1073/pnas.1417138111 (2014).

62 Patel, D. S. et al. Dynamics and Interactions of OmpF and LPS: Influence on Pore Accessibility and Ion Permeability. Biophysical journal 110, 930–938, doi:10.1016/j.bpj.2016.01.002 (2016).

63 Kumar, R., Iyer, V. G. & Im, W. CHARMM-GUI: A graphical user interface for the CHARMM users. Abstr Pap Am Chem S 233, 273–273 (2007).

64 Jorgensen, W. L., Chandrasekhar, J., Madura, J. D., Impey, R. W. & Klein, M. L. Comparison of Simple Potential Functions for Simulating Liquid Water. J Chem Phys 79, 926–935, doi:Doi 10.1063/1.445869 (1983).

65 Phillips, J. C. et al. Scalable molecular dynamics with NAMD. J Comput Chem 26, 1781–1802, doi:10.1002/jcc.20289 (2005).

66 Huang, J. et al. CHARMM36m: an improved force field for folded and intrinsically disordered proteins. Nature methods 14, 71–73, doi:10.1038/nmeth.4067 (2017).

67 Guvench, O. et al. Additive Empirical Force Field for Hexopyranose Monosaccharides. J Comput Chem 29, 2543–2564, doi:10.1002/jcc.21004 (2008).

68 Klauda, J. B. et al. Update of the CHARMM All-Atom Additive Force Field for Lipids: Validation on Six Lipid Types. J Phys Chem B 114, 7830–7843, doi:10.1021/jp101759q (2010).

69 Jo, S. et al. Lipopolysaccharide membrane building and simulation. Methods in molecular biology 1273, 391–406, doi:10.1007/978-1-4939-2343-4_24 (2015).

70 Shaw, D. E. et al. Anton 2: Raising the bar for performance and programmability in a special-purpose molecular dynamics supercomputer. Int Conf High Perfor, 41–53, doi:10.1109/Sc.2014.9 (2014).

71 Lippert, R. A. et al. Accurate and efficient integration for molecular dynamics simulations at constant temperature and pressure. J Chem Phys 139, doi:Artn 164106 10.1063/1.4825247 (2013).

72 Ritchie, T. et al. Reconstitution of membrane proteins in phospholipid bilayer nanodiscs. Methods in enzymology 464, 211–231 (2009).

