## Supplementary Information for "Lipoprotein sorting to the cell surface via a crosstalk between the Lpt and Lol pathways during outer membrane biogenesis"

### Extended Data Figure 1

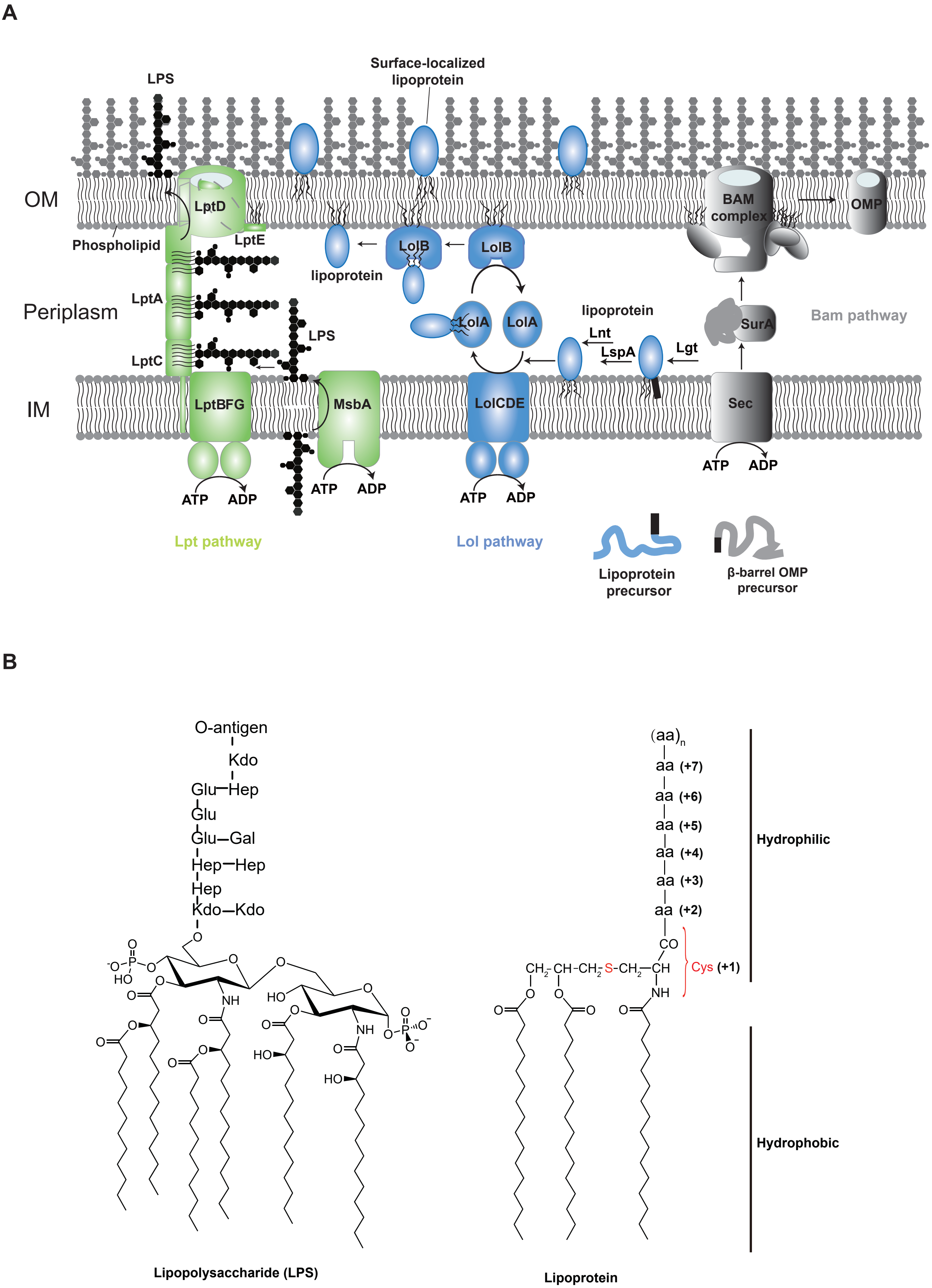

### Extended Data Figure 2

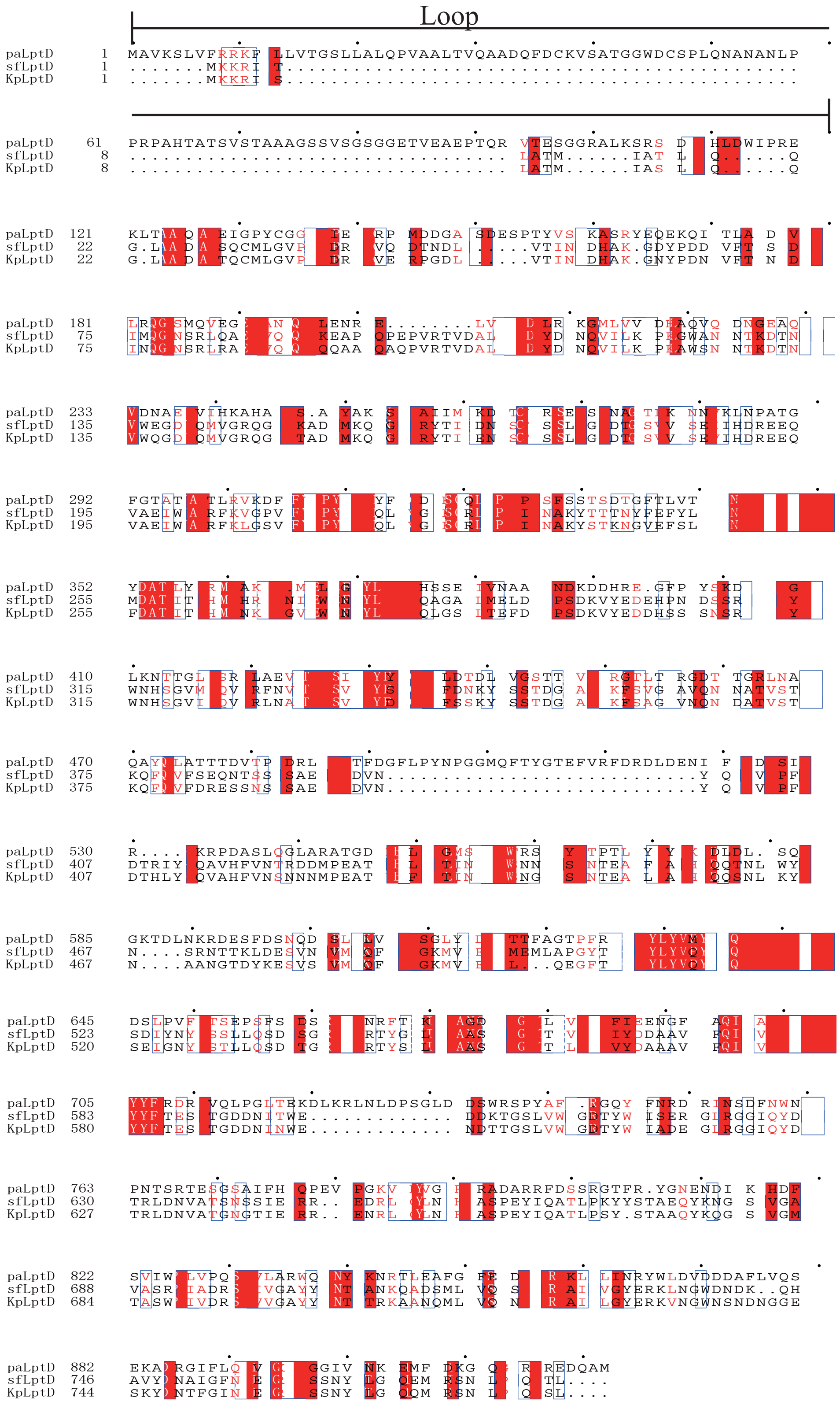

#### Extended Data Figure 3

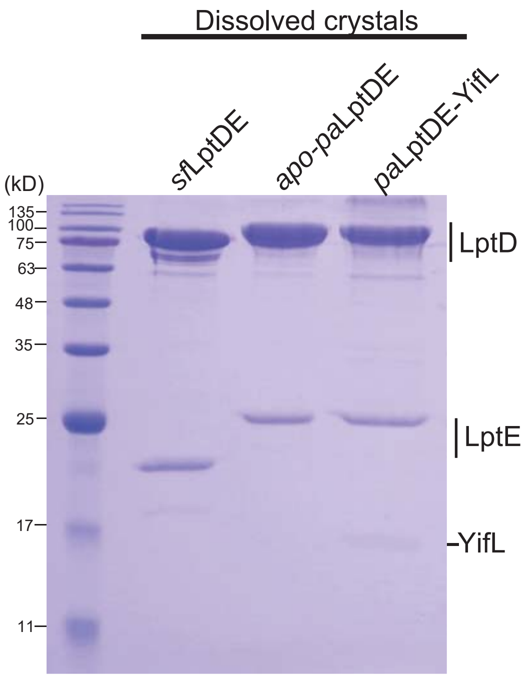

### Extended Data Figure 4

**A**

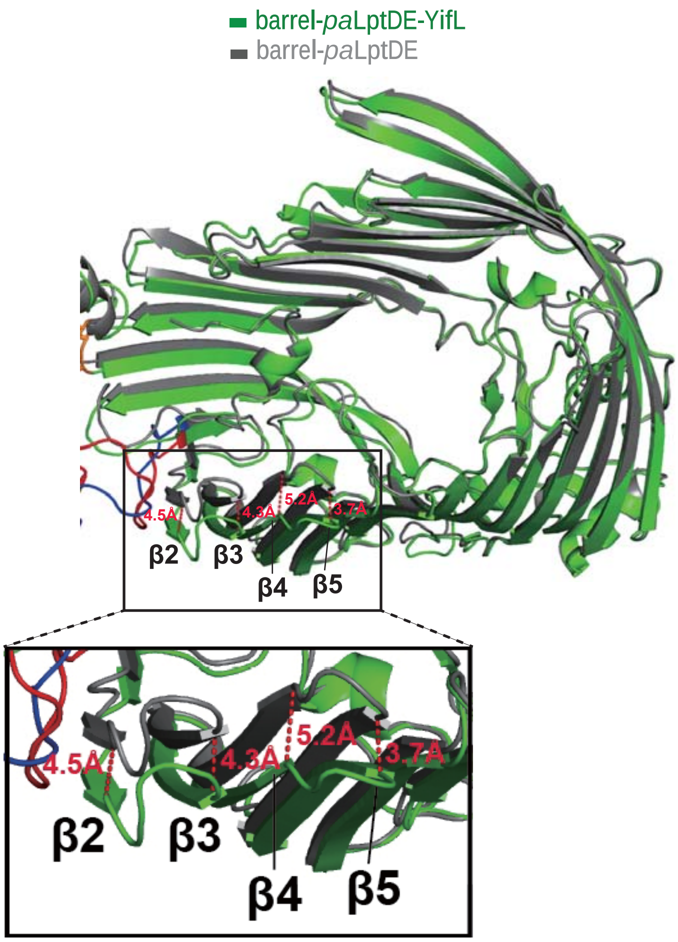


**B**

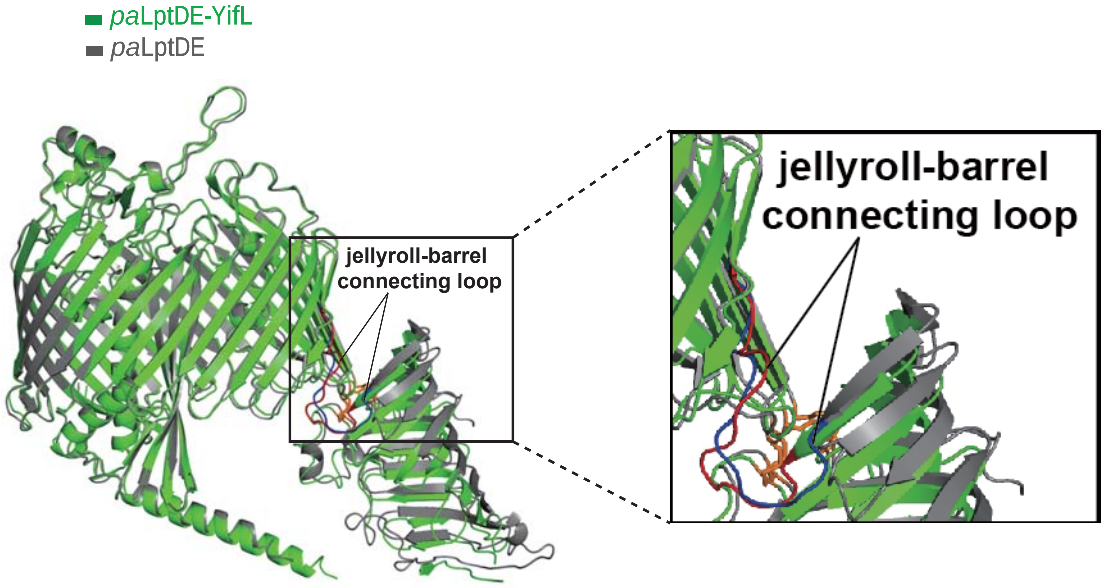

**Extended Data Figure 5**

A

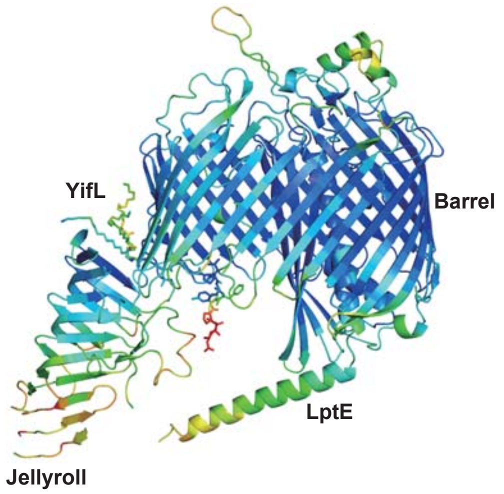


*pa*LptDE-YifL

B

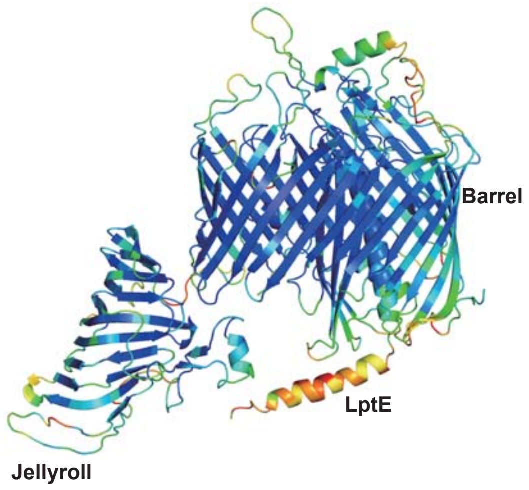


apo-*pa*LptDE

### Extended Data Figure 6

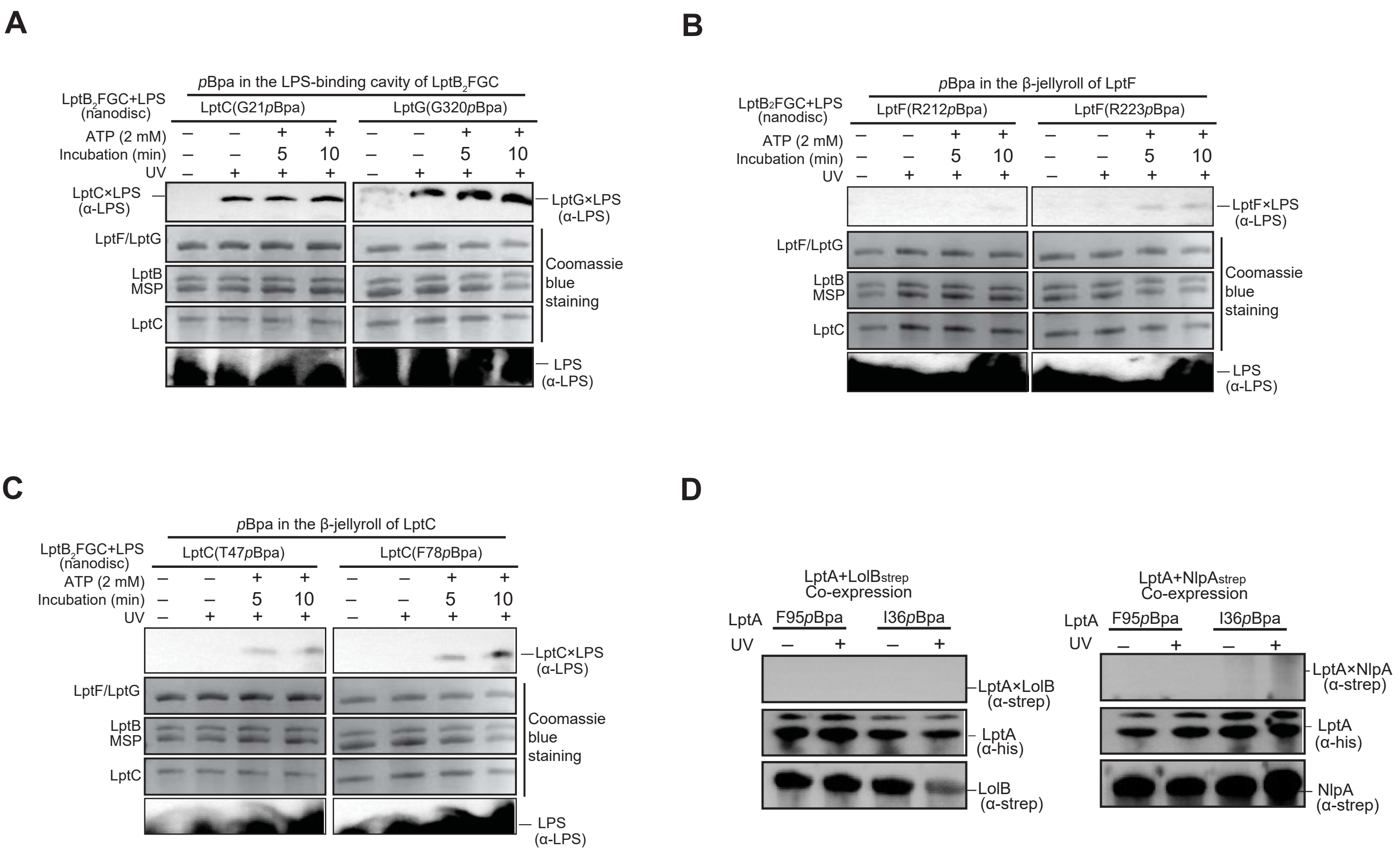

### Extended Data Figure 7

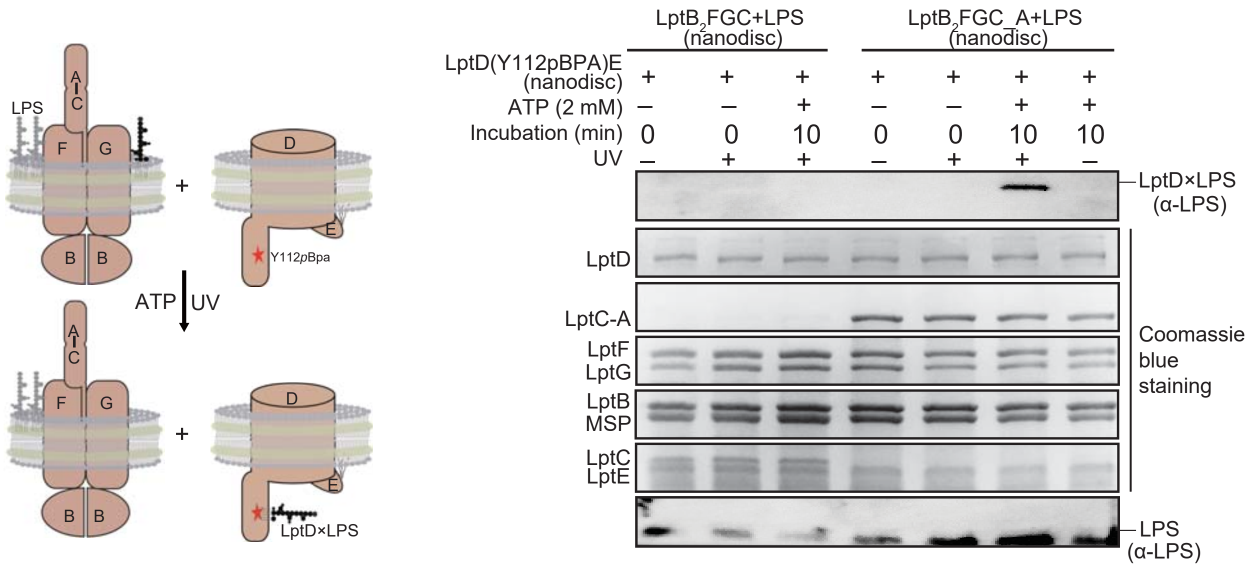

**Extended Data Figure Legends**

**Extended Data Fig. 1. Three established pathways responsible for OM biogenesis in Gram-negative bacteria.** (**A**) The Lpt, Lol and Bam pathways are responsible for transport of LPS, lipoproteins and β-barrel OMPs to the OM, respectively. Three membrane-bound enzymes Lgt, LspA and Lnt catalyze lipoprotein maturation upon being translocated across the IM by the SecYEG translocon. Lipoproteins sort to the inner leaflet of the OM via the Lol pathway. Surface-localized lipoprotein (SLP) anchors to the outer leaflet of the OM, with its C-terminus exposed in extracellular milieu. LPS transport to the cell surface via the Lpt pathway requires the formation of a transenvelope complex that consists of seven essential Lpt proteins, LptA-G. (**B**) Chemical structures of LPS (six acyl chains) and lipoproteins (three acyl chains). Both LPS and lipoproteins consists of hydrophobic acyl chains and hydrophilic sugar/polypeptide moiety. The first residue of a matured bacterial lipoprotein a Cys residue and that was covalently linked with three acyl chains.

**Extended Data Fig. 2. Amino acid sequence alignment for LptD homologs from different bacterial strains.** *Pa*LptD contains an approximately 120-residue Loop, and it is labeled above the sequence. *Pseudomonas aeruginosa* LptD, *pa*LptD*;* *Shigella flexneri* LptD, *sf*LptD; and *Klebsiella pneumonia* LptD, *kp*LptD. Alignment was made using Clustal O and colored in ESPript.

**Extended Data Fig. 3. SDS-PAGE analysis of dissolved crystals.** The apo-*pa*LptDE crystals and the *pa*LptDE-YifL crystals were obtained by using proteins purified from the *yifl*-depleted *E. coli* strain SF100 [KS272Δ (*ompT-entF*)] and *E. coli* strain SF100 [KS272Δ (*ompT-entF*)], respectively. The *sf*LptDE crystals that were obtained by using proteins purified from *E. coli* strain SF100 [KS272Δ (*ompT-entF*)] did not contain noticeable YifL. The SDS-PAGE gel was stained by coomassie blue.

**Extended Data Fig. 4. Structural overlay of *pa*LptDE with *pa*LptDE-YifL.** (**A**) Overlay of the β-barrel domain of *pa*LptD in *pa*LptDE (grey) with that in *pa*LptDE-YifL (green) showing the enlarged β-barrel (Strands β2, β3, β4 and β5) in *pa*LptDE-YifL on the periplasmic side. For clarity, the β-jellyroll domain of *pa*LptD, YifL and *pa*LptE are not shown. (**B**) Structural overlay showing a significant rearrangement of the jellyroll-barrel connecting loop between *pa*LptDE (grey) and *pa*LptDE-YifL (green). The jellyroll-barrel connecting loops are highlighted in red (*pa*LptDE-YifL) and blue (*pa*LptDE), respectively. For clarity, YifL is omitted.

**Extended Data Fig. 5. Comparison of crystallographic B factors of different domains (and LptE) in *pa*LptDE-YifL and apo-*pa*LptDE structures.** (**A-B**) Crystallographic B factors (Cα atoms) colors the structures of *pa*LptDE-YifL (**A**) and *pa*LptDE(**B**) in PyMOL with the blue_white_red palette. The average B factors (Cα atoms) of individual domains of *pa*LptDE-YifL: β-jellyroll (110.5 $Å^2$), β-barrel (59.6 $Å^2$), LptE(54.2 $Å^2$) and YifL(64.6 $Å^2$); The average B factors (Cα atoms) of individual domains of *pa*LptDE: β-jellyroll (95.3 $Å^2$), β-barrel (62.3 $Å^2$) and LptE (50.3 $Å^2$).

**Extended Data Fig. 6. *In vitro* $LptB_2FGC$-mediated LPS binding and transport assay (A-C).** (**A**) LptC (G21*p*Bpa) and LptG (G320*p*Bpa) are in the LPS-binding cavity of $LptB_2FGC$; (**B**) LptF (R212*p*Bpa) and LptF (R223*p*Bpa) are in the β-jellyroll domain of LptF; (**C**) LptC (T47*p*Bpa) and LptC (F78*p*Bpa) are in the β-jellyroll domain of LptC. All the *p*Bpa-substituted sites in $LptB_2FGC$ cross-linked to LPS that was reconstituted in nanodiscs together with $LptB_2FGC$ upon UV radiation. LPS cross-linked to *p*Bpa substituted sites in the β-jellyroll domains of LptF and LptC are in an ATP-dependent manner. LPS antibody was used for detection of the crosslinks. (**D**) Neither LolB nor NlpA cross-linked to $LptA^{F95pBpa}$ or $LptA^{I36pBpa}$ when co-expressed in *E. coli*. For immunoblotting detection, LptA and LolB/NlpA contained a C-terminal His and Strep tag II, respectively.

**Extended Data Fig. 7. *In vitro* reconstitution of membrane-to-membrane LPS transport.** Both $LptB_2FGC$-A (or $LptB_2FGC$) and LPS were reconstituted into nanodiscs. After being incubated with nanodisc-embedded $LptD^{Y112pBpa}E$, photocrosslinking was performed. The LPS cross-linked to $LptD^{Y112pBpa}$ in an ATP-dependent manner, and LptA is required for LPS transport to LptD. LPS antibody was used for detection of the crosslinks. Left cartoons show experimental designs of the LPS transport assay. The red star denotes the *p*Bpa site in the β-jellyroll domain of LptD.

**Extended Data Table 1: *pa*LptDE-bound lipoproteins identified by MS**

| Protein names | Accession No. | Residues at positions +1 - +8 | Total residues (full-length) | Mol. mass of mature lipoprotein (M.W. of fatty acyl chains=818 Da) |
|---|---|---|---|---|
| **YifL** | AJE58371.1 | CGLKGPLY | 67 | 5916.6 |
| **Lpp** | AUO32430.1 | CGQKGPLY | 78 | 7203.0 |
| **YedD** | ART43778.1 | CAEVENYN | 137 | 14366.4 |
| **YbjP** | APQ22223.1 | CTTVTPAY | 171 | 17802.8 |
| **SlyB** | APQ21448.1 | CVNNDTLS | 155 | 14636.4 |
| (**OipY**) | APQ19979.1 | CVAAAVVG | 191 | 19025.6 |
| **Slp** | APQ21285.1 | CVTVPDAI | 193 | 19494.3 |
| **Putative lipoprotein** | APQ22800.1 | CAKPPTTI | 192 | 19846.5 |
| **Blc** | APQ23470.1 | CSSPTPPR | 177 | 18860.3 |

**Extended Data Table 2: Theoretical and experimental masses of LptDE associated with lipoproteins**

| Complexes | Theoretical mass (Da) (with molecular mass of fatty acyl chains, 818 Da) | Experimental mass (Da) |
|---|---|---|
| *sf*LptDE$_{6\times His}$ | 108,162 | 108,172 |
| *sf*LptDE$_{6\times His}$+YifL | 114,079 | 114,089 |
| *sf*LptDE$_{6\times His}$+YedD | 122,528 | 122,518 |
| *pa*LptD$^{\Delta 12}$E$_{6\times His}$ | 123,481 | 123,502 |
| *pa*LptD$^{\Delta 12}$E$_{6\times His}$+YifL | 129,398 | 129,418 |
| *pa*LptD$^{\Delta 23}$E$_{6\times His}$ | 121,162 | 121,180 |
| *pa*LptD$^{\Delta 23}$E$_{6\times His}$+YifL | 127,078 | 127,052 |
| *pa*LptD$^{\Delta 23}$E$_{6\times His}$+YbjP | 138,965 | 138,950 |
| *pa*LptD$^{\Delta 23}$E$_{6\times His}$+SlyB | 135,798 | 135,838 |

Note: *pa*LptD$^{\Delta 12}$and *pa*LptD$^{\Delta 23}$ denote the *pa*LptD variants that the first 12 and 23 residues of the matured *pa*LptDE were degraded.

**Extended Data Table 3: Distance change between two ends of each strand in the β-jellyroll of *pa*LptD with or without YifL binding**

| No. of jellyroll β strands | *pa*LptDE (Å) | *pa*LptDE-YifL (Å) | Change of distance (Å) |
|---|---|---|---|
| **β11** (F278-P289) | 25.7 | 22.8 | **2.9** |
| **β10** (G265-V275) | 28.2 | 23.9 | **4.3** |
| **β9** (W250-N260) | 25.1 | 21.2 | **4.1** |
| **β8** (A231-T241) | 23.2 | 20.3 | **2.9** |
| **β7** (A218-S228) | 24.1 | 20.2 | **3.9** |
| **β6** (A201-I213) | 19.1 | 17.0 | **2.1** |
| **β5** (V197-M188) | 18.1 | 16.8 | **1.3** |
| **β4** (R174-D185) | 20.6 | 18.8 | **1.8** |
| **β3** (M159-H169) | 19.2 | 18.7 | **0.5** |
| **β2** (A146-Q156) | 19.3 | 16.6 | **2.7** |
| **β1** (Y131-Y139) | 19.9 | 16.3 | **3.6** |
